# Glycerol metabolism triggers trypanosome differentiation into transmissible forms in mammalian tissue-like conditions

**DOI:** 10.1101/2025.05.26.656071

**Authors:** Mohammad El Kadri, Fabien Guegan, Parul Sharma, Estefania Calvo Alvarez, Erika Pineda, Pauline Morand, Jean Marc Tsagmo Ngoune, Justine Escard, Jean-William Dupuy, Aline Rimoldi, Nicolas Plazolles, Marc Biran, Michael P. Barrett, Luisa M. Figueiredo, Brice Rotureau, Frédéric Bringaud

**Affiliations:** Univ. Bordeaux, CNRS, Microbiologie Fondamentale et Pathogénicité (MFP), UMR 5234, F-33000 Bordeaux, France; Gulbenkian Institute for Molecular Medicine, 1649-028 Lisbon, Portugal; Trypanosome Transmission Group, Trypanosome Cell Biology Unit, Institut Pasteur, Université Paris Cité, INSERM U1347, Paris, France; Sorbonne Université, ED515 Complexité du Vivant, Paris, France; Centre de Génomique Fonctionnelle, Plateforme Protéome, Université de Bordeaux, 146 rue Léo Saignat, 33076 Bordeaux, France; Univ. Bordeaux, CNRS, INSERM, TBM-Core, US5, UAR 3427, OncoProt, F-33000 Bordeaux, France; Univ. Bordeaux, CNRS, Centre de Résonance Magnétique des Systèmes Biologiques (CRMSB), UMR 5536, F-33000 Bordeaux, France; Wellcome Centre for Integrative Parasitology, Institute of Infection, Immunity and Inflammation, College of Medical, Veterinary and Life Sciences, University of Glasgow, Glasgow, United Kingdom; Glasgow Polyomics, Wolfson Wohl Cancer Research Centre, Garscube Campus, College of Medical Veterinary and Life Sciences, University of Glasgow, Glasgow, United Kingdom; Parasitology Unit, Institut Pasteur of Guinea, Conakry, Guinea

## Abstract

In the mammalian bloodstream, *Trypanosoma brucei*, the parasite responsible for sleeping sickness, proliferates as slender forms before undergoing quorum-sensing (QS)-mediated differentiation into cell cycle-arrested stumpy forms (stumpy-QS), a transition that regulates parasitaemia and primes parasites for tsetse fly transmission. Beyond the bloodstream, *T. brucei* also occupies extravascular, adipose-rich tissues such as the skin, a potential reservoir for transmission. Here, we identify an alternative slender-to-stumpy differentiation pathway driven by glycerol, a host metabolite abundant in adipose-rich tissues, that could resolve the long-standing paradox of successful parasite transmission from human hosts during chronic infection despite low parasitaemia. We show that high (10 mM) and non-physiological glycerol concentrations under low glucose conditions (0.5 mM) induce differentiation, generating distinct stumpy-Glyc forms that resemble stumpy-QS parasites but have an extended lifespan. Under conditions mimicking dermal tissue interstitial fluids (4 mM glucose, 0.25 mM glycerol), we show that glycerol promotes the emergence of proliferative intermediate forms that retain transmission potential and can differentiate into fly host-specific procyclic forms *in vitro* and within tsetse flies. These findings open the door for reevaluation of the model of *T. brucei* transmission and supports a dominant role for adipocyte-derived glycerol in the skin in sustaining parasite transmission.

## Introduction

*Trypanosoma brucei* is an extracellular parasite that causes the neglected tropical disease human African trypanosomiasis (HAT), also known as sleeping sickness, and animal African trypanosomiases (AAT), such as *nagana*. This parasite undergoes a complex life cycle as it transits from the bloodstream of a mammalian host in bloodstream forms (BSF) to the alimentary tract in procyclic forms (PCF) and salivary glands of its blood-feeding insect vector, the tsetse fly. The extracellular fluids of the mammalian host, including blood and interstitial fluids, are rich in glucose, the main carbon source used by BSF trypanosomes to fuel their central carbon metabolism and ATP production. However, glucose can be successfully replaced by glycerol *in vitro* to feed the parasite’s central metabolism, suggesting that this alternative carbon source may also play a role in trypanosome infection *in vivo* [1–3]. Indeed, recent studies showed that BSF trypanosomes reside primarily in the extravascular space of tissues such as skin [4,5] and adipose tissue [6], where interstitial glycerol concentrations are 5 to 20 folds higher than in plasma [7–8]. Glycerol is released from adipocytes by both lipolysis and lipolysis-independent processes such as glycolysis [9,10]. It has also recently been reported that trypanosome-induced adipocyte lipolysis may protect the host against escalation of trypanosome infection, by releasing toxic amounts of fatty acids [11]. Taken together, these data suggest that adipocyte-trypanosome metabolic interactions, possibly through glycerol, may play an important role in trypanosome infection.

During ascending development of the parasites in the blood of the mammalian hosts (parasitaemia), rapidly dividing BSF with a slender morphology (slender forms) predominate in blood and tissues. At the peak of a parasitaemic wave, slender forms differentiate into shorter so-called stumpy forms, which are irreversibly growth-arrested in G_0_/G_1_ phase. These arrested stumpy parasites have a limited lifespan in the host blood. Slender-to-stumpy differentiation relies on a quorum sensing mechanism that is triggered by the accumulation of a ‘‘stumpy induction factor’’ (SIF) [12], which was recently identified as a mixture of di- and tripeptides produced by oligopeptidases excreted by the parasites [13,14]. Quorum sensing not only protects the host by preventing lethal high parasitaemia, but also results in the generation of stumpy forms that are pre-adapted to cyclical development, as they rapidly differentiate into PCF when ingested by tsetse flies during a blood meal, enabling transition to the next life stages of the parasite in the fly. However, a paradox has long persisted in the field: low parasitaemia is frequently observed in patients, but the triggering of quorum sensing for differentiation into stumpy forms is dependent on high parasite density [15]. Here, we show that, distinctly from the SIF-based quorum sensing mechanism, glycerol is sufficient to induce the differentiation of slender forms into long-lived, cell cycle-arrested stumpy forms, or proliferative intermediate forms, both capable of infecting the insect vector.

## Results

### Glycerol induces the differentiation of slender BSF into cell cycle-arrested stumpy forms

Across the transmission of trypanosomes from the mammalian host to the insect vector, the parasites differentiate first from the slender forms to the stumpy forms, distinguished by the expression of the cell surface protein PAD1 (carboxylate surface transporter family), in the mammalian host, and then from the stumpy form to the PCF form, distinguished by the cell surface protein EP-procyclin (procyclic acidic repetitive protein), in the fly [16,17]. These two successive differentiation events required for the transmission of trypanosomes from the mammalian host to the insect vector can be reproduced *in vitro*. First, differentiation from slender forms to growth-arrested forms, which have the morphological and molecular characteristics of stumpy forms, can be induced *in vitro* by (*i*) high parasite density, which leads to the excretion of large amounts of oligopeptidases involved in SIF production, (*ii*) addition of the SIF-rich brain heart infusion (BHI) medium [13], or (*iii*) addition of a cyclic AMP analogue (8-pCPT-cAMP), which bypasses the initial signaling events [12]. Second, stumpy-to-PCF differentiation can be induced *in vitro* by the addition of cis-aconitate and a temperature switch from 37°C to 27°C.

We first assessed the role of glycerol on slender BSF differentiation using the pleomorphic *T. brucei* AnTat1.1E-GFP::PAD1_UTR_, containing a *GFP* gene fused to the 3’ UTR of stumpy-form associated *PAD1* gene, which enables maximum expression in stumpy forms [6]. We compared the fate of the slender AnTat1.1E-GFP::PAD1_UTR_ incubated under five different conditions, four of which represent known differentiation inducing conditions outlined above and one providing glycerol in the presence of low glucose. These five conditions are: (*i*) high cell density (HCD) in glucose-rich (10 mM) Creek’s minimal medium [18] (CMM_Glc10/HCD), (*ii*) CMM_Glc10 containing 5% SIF-rich BHI medium [13] (CMM_Glc10/BHI), (*iii*) CMM_Glc10 containing 100 µM 8-pCPT-cAMP [12] (CMM_Glc10/CPT), (*iv*) SIF-rich CMM_Glc10 spent medium derived from a stationary phase culture of AnTat1.1E-GFP::PAD1_UTR_ (CMM_Glc10/SIF), and (*v*) low glucose (0.5 mM) CMM containing 10 mM glycerol (CMM_Glc0.5/Glyc10). We observed that the addition of 10 mM glycerol under low glucose conditions (CMM_Glc0.5/Glyc10) induced phenotypic effects similar to known conditions that induce stumpy-QS. This includes growth arrest (Fig.1a), further confirmed by reduced DNA synthesis as assessed by 5-ethynyl-2-deoxyuridine (EdU) incorporation (Figs.1b and S1) and an accumulation of cells in G1/G0/S phases accompanied by a decrease in dividing cells (G2/M & post-mitotic phases) (Fig.1c). Consistent with the transition to stumpy forms, this 10 mM glycerol under low glucose condition also induced an increase in GFP::PAD1_UTR_ and PAD1 expression, as assessed by cell cytometry (Fig.1d), Western blotting (Fig.1e) and immunofluorescence analysis (Fig.S2). Lastly, this condition also induced an increase in cell width and the distance between the nucleus and the kinetoplast (Figs.1f and S3), indicating progression towards a typical stumpy morphology. These features, characteristic of stumpy-QS forms, demonstrate that glycerol is sufficient to induce the differentiation from slender to stumpy-like forms, which we named stumpy-Glyc.

**Figure 1.**
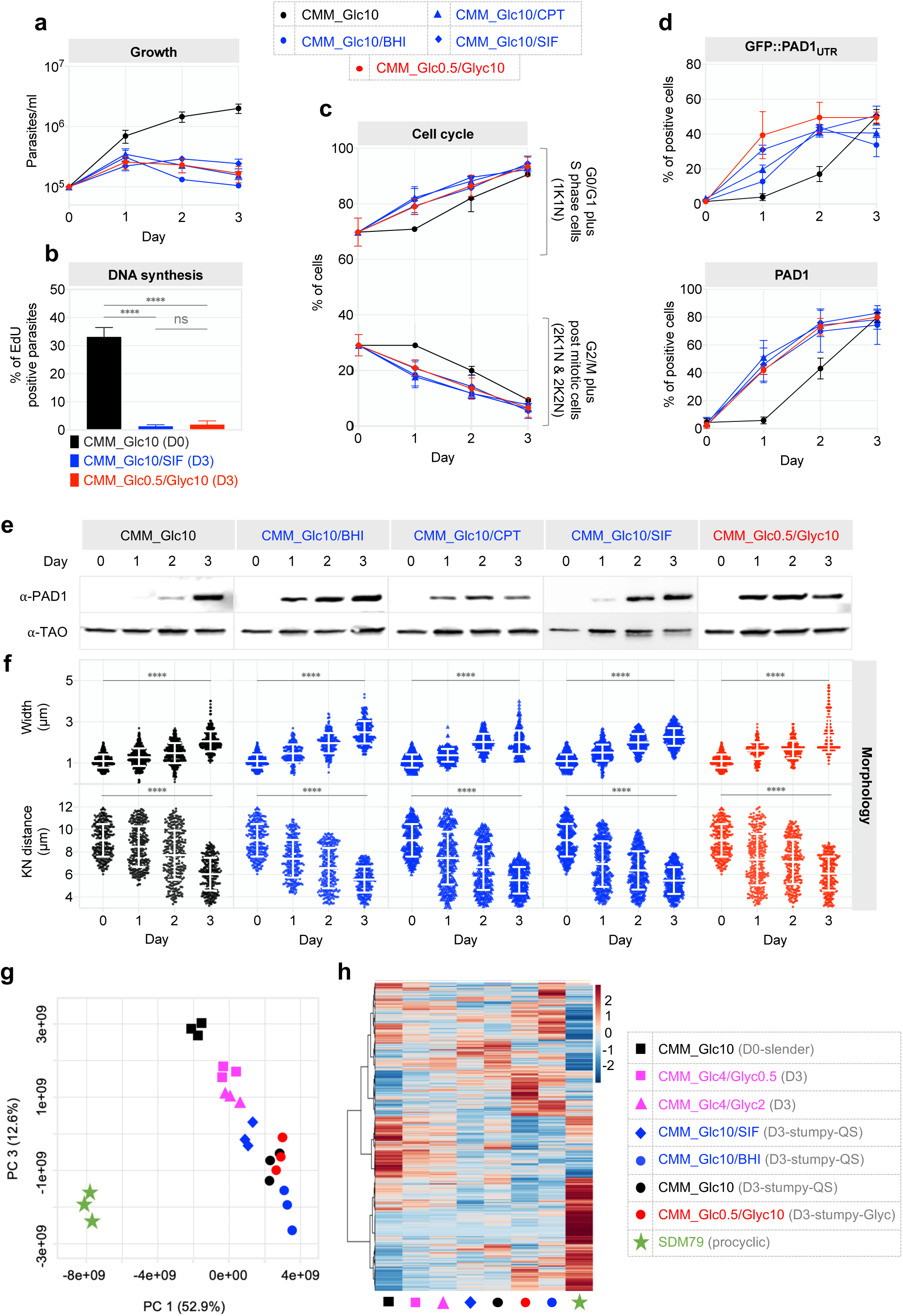
Glycerol induces production of stumpy-Glyc forms. **a**, Growth without dilution of the pleomorphic AnTat1.1E-GFP::PAD1_UTR_ strain incubated under five growth conditions leading to differentiation into stumpy-QS or stumpy_Glyc, *i.e.* (*i*) Creek’s minimal medium containing 10 mM glucose (CMM_Glc10), which allows production of stumpy-QS at high cell density, (*ii*) SIF-rich CMM_Glc10 produced from a stationary phase culture of AnTat1.1E-GFP::PAD1_utr_ (CMM_Glc10/SIF), (*iii*) CMM_Glc10 supplemented with 5% of SIF-rich brain-heart infusion broth (CMM_Glc10/BHI), (*iv*) CMM_Glc10 supplemented with 100 µM of cAMP analogue 8-pCPT-2’-O-Me-cAMP (CMM_Glc/CPT) and (*v*) CMM containing 0.5 mM glucose and 10 mM glycerol (CMM_Glc0.5/Glyc10). y-axis shows parasites per ml on a log10 scale**. b**, Parasite proliferation was assessed from microscopy images of EdU labelled (*i*) slender forms incubated in CMM_Glc10 and collected at low cell density (here is represented by day 0), (*ii*) stumpy-QS forms derived from a 3 days culture in (CMM_Glc10/SIF) and (*iii*) stumpy-Glyc derived from a 3 days culture in (CMM_Glc0.5/Glyc10). More than 300 cells per group, (****, p < 0.0001; ns, not significant). **c**, the top panel shows the percentage of 1K1N cells, with 1 kinetoplast (K; mitochondrial DNA) 1 nucleus (N), which correspond to G0/G1 and S-phase cells, while the bottom panel shows the percentage of dividing cells, including 2K1N (G2/M-phase cells) and 2K2N (post-mitotic cells). More than 300 cells per group. **d**, Time course for the expression of the GFP::PAD1_UTR_ (top panel) and PAD1 using the ⍺-PAD1 immune serum (bottom panel) monitored by flow cytometry. **e**, Expression of PAD1 was confirmed by western blot (⍺-PAD1) using the ⍺-TAO immune serum as a loading control. **f**, Stumpy morphology was confirmed by measuring the cell width and the distance between the kinetoplast and the nucleus (K-N) using a previously described ImageJ macro. More than 300 cells per group, (****, p < 0.0001). The proteomic data described in figures (Fig.S4-S7) are represented by the principal component analysis (**g**) and the heat map (**h**) encompassing all the 5,464 quantified proteins identified with at least 2 unique peptides. This illustrates major proteome remodeling between slender (or stumpy-Glyc) maintained at low cell density in CMM_Glc10 (CMM_Glc10/LCD), CMM_Glc4/Glyc0.5, CMM_Glc4/Glyc4 and CMM_Glc0.5/Glyc10, stumpy-QS forms (CMM_Glc10/SIF, CMM_Glc10/BHI and CMM_Glc10/LCD) and freshly *in vitro* differentiated PCF (SDM79). Data are presented as the mean ±SD of three independent replicates. Error bars smaller than symbols are not displayed for clarity.

To compare the stumpy-Glyc forms with stumpy forms induced by known differentiation conditions and with other trypanosome forms, we examined the proteome of stumpy-Glyc, stumpy-QS forms produced under CMM_Glc10/HCD, CMM_Glc10/BHI and CMM_Glc10/SIF conditions, and the proteome of PCF and slender BSF. Principal component analysis across these proteomes showed that the stumpy-QS and stumpy-Glyc cells clustered together, far away from cells in the slender and PCF forms (Fig.1g). The heat map encompassing the 5,464 z-scored LFQ-quantified protein groups (all with at least two peptides) showed that there is major proteome remodeling between slender, stumpy and procyclic forms (Fig.1h). Among these 5,464 proteins, 109 and 59 were significantly up-and down-regulated, respectively, in the stumpy-Glyc compared to the slender BSF (Fig.S4). Similarly, there was significant regulation of 110, 139, 73 (up-regulated) and 80, 99, 64 (down-regulated) proteins in stumpy-QS forms as induced by CMM_Glc10/HCD, CMM_Glc10/BHI and CMM_Glc10/SIF, respectively, compared to slender BSF (Fig.S4). Consistent with the proximity of the different types of stumpy forms, 37 and 12 of the significantly up- and down-regulated proteins compared to slender BSF, respectively, were shared between stumpy-Glyc and stumpy-QS obtained in three different ways (CMM_Glc10/HCD, CMM_Glc10/BHI and CMM_Glc10/SIF) (Figs.S5a and S5e). In addition, GO term enrichment analysis for biological process of differentially expressed proteins showed equivalent profiles for stumpy-Glyc and stumpy-QS obtained in three same different ways (Fig.S6). These data were confirmed by Western blot analyses for seven proteins up-regulated during the slender-to-stumpy transition induced by glycerol, HCD, BHI and SIF (Fig.S7). This is consistent with previous observations for stumpy-QS [19,20].

### Glycerol-induced Stumpy-Glyc forms have a longer lifespan than stumpy-QS forms

To investigate whether the glycerol-induced differentiation of slender into stumpy-Glyc forms relies on a cell density-dependent quorum sensing mechanism, slender cells were cultured under various growth conditions starting from different initial inoculum densities (Fig.2). Under CMM_Glc10 conditions, the differentiation to stumpy-QS forms occurred through a cell density-dependent quorum-sensing mechanism as GFP::PAD1_UTR_ expression peaks only when cells reach high densities (Fig.2a). In contrast, glycerol immediately triggered GFP::PAD1_UTR_ expression, regardless of the initial inoculum density, and was accompanied by immediate growth arrest within one day (Fig.2b). In addition, while stumpy-QS cells died a few days after reaching high cell densities, stumpy-Glyc cells remained alive for at least 8 days and maintained a high level of GFP::PAD1_UTR_ expression.

**Figure 2.**
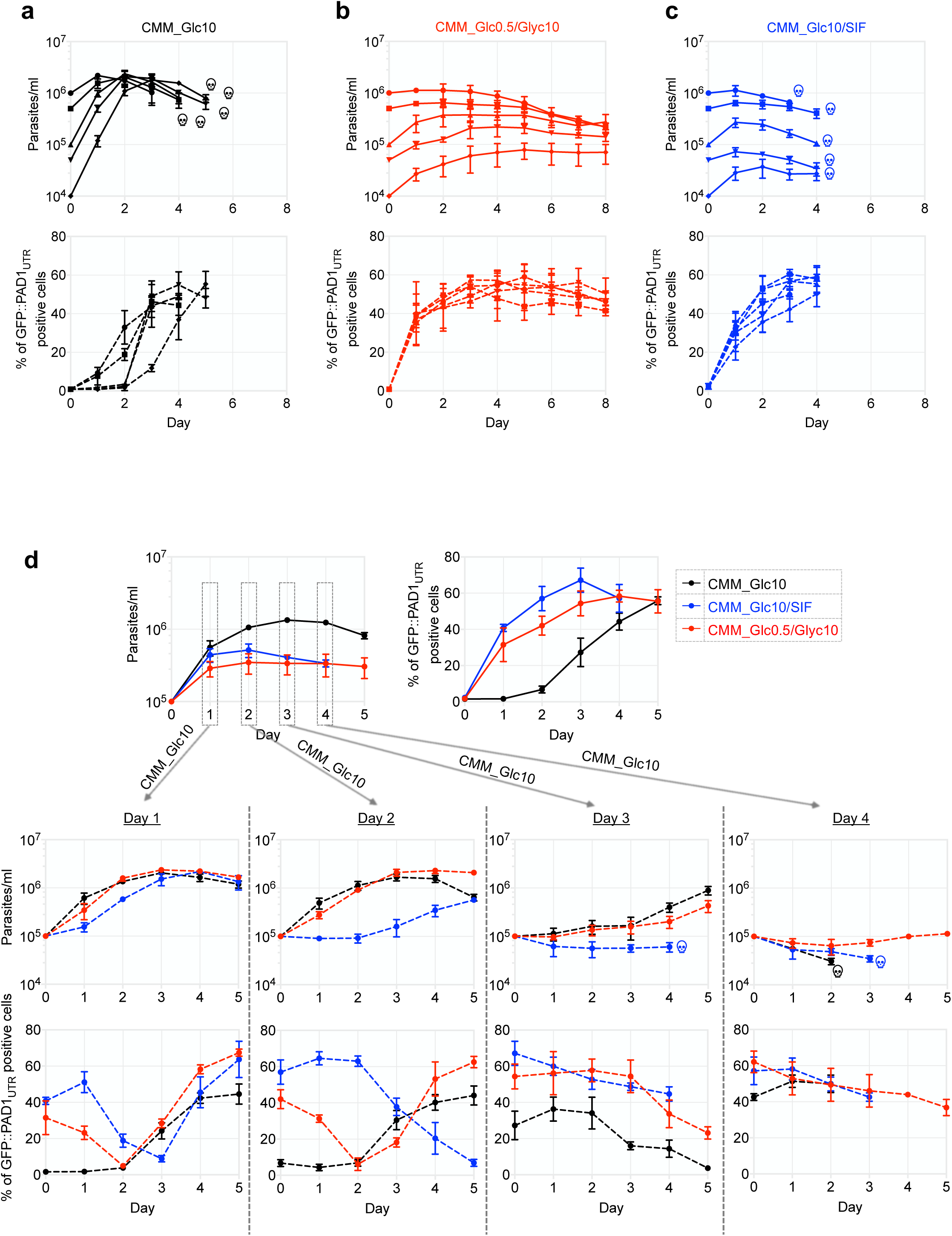
Stumpy-Glyc are produced under high density-independent conditions and persist for longer than stumpy-QS. Growth of the AnTat1.1E-GFP::PAD1_UTR_ strain, without dilutions, in CMM_Glc10 (**a**), CMM_Glc0.5/Glyc10 (**b**) and CMM_Glc10/SIF (**c**) from different initial inoculum cell densities (10⁴, 5 × 10⁴, 10⁵, 5 × 10⁵, and 10⁶ cells/ml) (top panel). The percentage of GFP::PAD1_UTR_ positive cells was determined every day by cell cytometry (lower panel). The skull symbolises the death of the entire cell population, which is defined microscopically by the loss of motility followed by the loss of cell integrity the next day. **d**, AnTat1.1E-GFP::PAD1_UTR_ slender forms, initially incubated under CMM_Glc10, CMM_Glc10/SIF, and CMM_Glc0.5/Glyc10 conditions (top panel), were daily washed and transferred to fresh CMM_Glc10 to assess their ability to revert and propagate as slender forms. The central panel illustrates continuous growth after transfer to CMM_Glc10, while the lower panel depicts GFP::PAD1_UTR_ expression. For all the growth curves, the y-axis shows parasites per ml on a log10 scale. Data are presented as the mean ±SD of three independent replicates. Error bars smaller than symbols are not displayed for clarity.

To examine whether the short lifespan of stumpy-QS was linked to stress induced by the high-density stationary phase, we repeated the experiment but under CMM_Glc10/SIF conditions so as to induce differentiation to stumpy-QS at low cell density, instead of high cell density. The addition of SIF to low-density slender parasites in culture immediately induced GFP::PAD1_UTR_ expression and growth arrest, but the stumpy-QS cells still proceeded to cell death 3 to 4 days later (Fig.2c), indicating that the death is density-independent and confirming the previously reported short lifespan of stumpy-QS *in vitro* of 3-4 days [21].

To compare the dynamics of growth arrest between stumpy-QS and stumpy-Glyc, AnTat1.1E-GFP::PAD1_UTR_ slender forms incubated under CMM_Glc10, CMM_Glc10/SIF and CMM_Glc0.5/Glyc10 conditions were washed and transferred to fresh CMM_Glc10 each of the 4 days after induction to access their potential ability to revert to and proliferate as slender forms. After one day of induction, the effects of glycerol (CMM_Glc0.5/Glyc10) and SIF (CMM_Glc10/SIF) were similar: no delay was observed before either stumpy-QS or stumpy-Glyc cells resumed proliferation (Fig.2d). At day 2, however, there was a two-day delay before CMM_Glc10/SIF-induced cells resumed proliferation, whereas this was unaffected for CMM_Glc0.5/Glyc10-induced cells (Fig.2d). This suggests that differentiation in stumpy-QS is stably committed earlier than differentiation in stumpy-Glyc. By day 4, both stumpy-QS (CMM_Glc10/SIF and CMM_Glc10/HCD) and stumpy-Glyc cells were no longer able to resume proliferation. As expected, stumpy-QS cells underwent cell death 2-3 days after transfer to fresh CMM_Glc10, whereas stumpy-Glyc remained alive until the end of the experiment on day 5 (Fig.2d).

### Glycerol must be metabolized to induce slender to stumpy-Glyc differentiation

Like glucose, glycerol is metabolized in the glycosomes and the cytosol, where it is mainly converted into pyruvate as an excreted end-product. To investigate the role of glycerol metabolism in the differentiation process, we generated individual mutant cell lines by deleting the *aquaglyceroporin* genes (*AQP1*, *AQP2* and *AQP3*), which are the only known proteins responsible for glycerol import, or by inactivating the five tandemly-arranged *glycerol kinase* genes (*GK*), which are responsible for glycerol conversion into glycerol 3-phosphate (Fig.3a-b). We observed that glycerol was no longer metabolized in the Δ*gk* and Δ*aqp1-3* cell lines, as indicated by the abolition of ^13^C-enriched pyruvate production from the metabolism of uniformly ^13^C-enriched glycerol ([U-^13^C]-glycerol) (Figs.3c and S8). Glycerol metabolism was restored by reintroduction of the *GK* and *AQP1* genes in the Δ*gk* (Δ*gk*/GK) and Δ*aqp1-3* (Δ*aqp1-3*/AQP1) mutants, respectively (Fig.3a-c). The Δ*gk* and Δ*aqp1-3* slender BSF mutants were unable to differentiate into stumpy-Glyc under CMM_Glc0.5/Glyc10 conditions, while retaining their ability to differentiate into stumpy-QS in the presence of SIF, as shown by monitoring cell growth and PAD1 expression (Figs.3d-e and S9). Differentiation to stumpy-Glyc was restored in the Δ*gk*/GK and Δ*aqp1-3*/AQP1 rescue cell lines (Fig.3d-e). These data demonstrated that glycerol must be metabolized at least up to glycerol 3-phosphate for slender parasites to differentiate into stumpy-Glyc forms and that stumpy-Glyc differentiation is mechanistically distinct from stumpy-QS differentiation, as stumpy-QS formation was not impacted in the *AQP* and *GK* null background.

**Figure 3.**
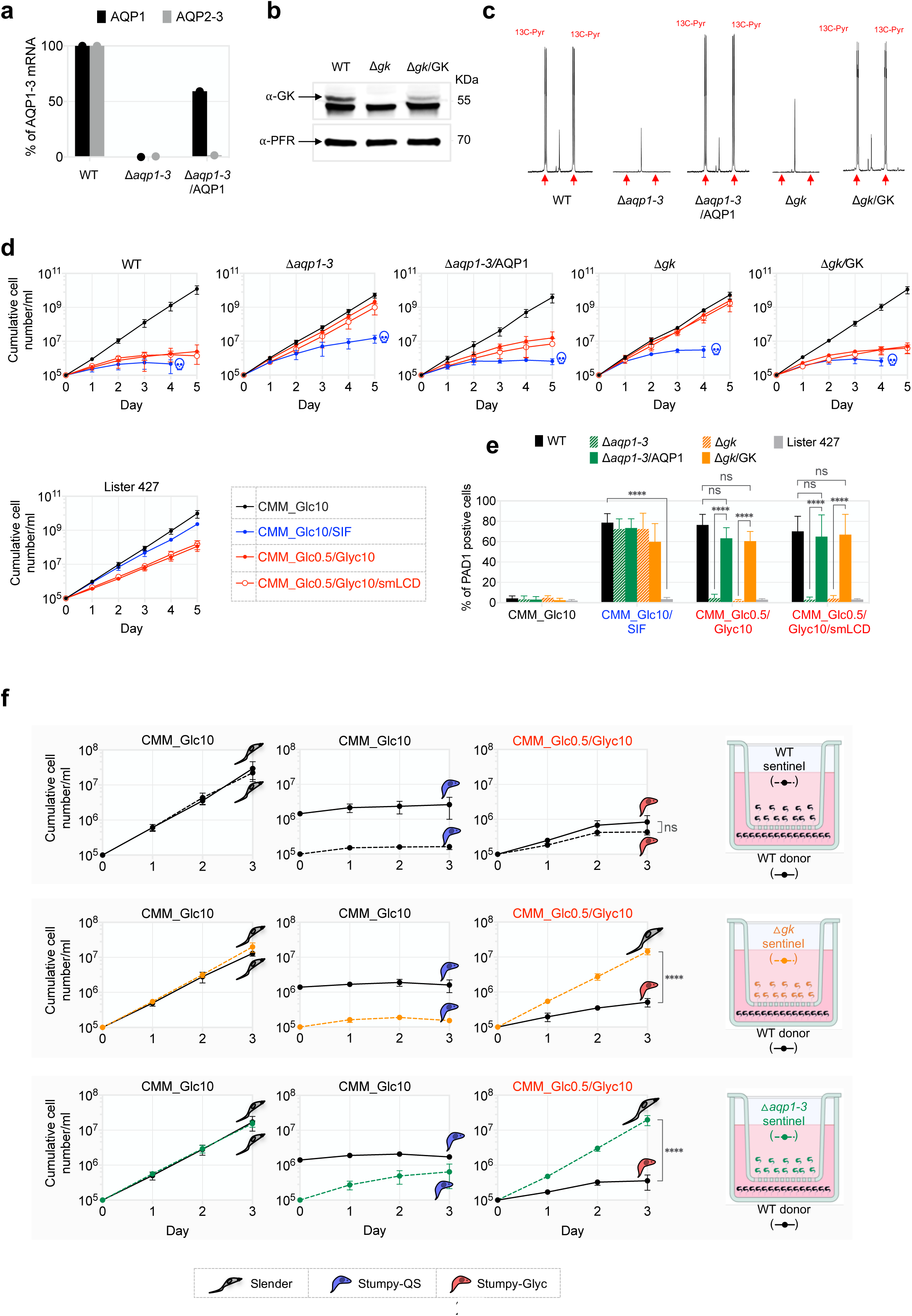
Glycerol metabolism is essential for differentiation and doesn’t induce the excretion of SIF-like factors. **a**, Expression of aquaglyceroporin (*AQP1* and *AQP2-3*) mRNA assessed by RT-qPCR by the comparative ΔΔC_T_ method of the WT (AnTat1.1E parental cell line), the AQP null (△*aqp1-3*) and the AQP1 rescued (△*aqp1-3*/AQP1) cell lines. **b**, Western blot analysis of the WT, glycerol kinase null mutant (△*gk*) and rescue (△*gk*/GK) cell lines with the anti-GK and the anti-PFR as a loading control. **c,** ^1^H-NMR-based analyses of the exometabolome validated the mutants cell line genotype. The parental, knockout and rescue cell lines were incubated for 1 h in PBS/NaHCO_3_ buffer containing 4 mM of uniformly ^13^C-enriched glycerol ([U-^13^C]-glycerol) before analysis of the spent medium by ^1^H-NMR (Bringaud et al.,2015). A part of each spectrum ranging from 2.0 and 2.6 ppm, corresponding to the pyruvate resonance, is shown. Resonances corresponding to ^13^C-enriched pyruvate produced from the metabolism of [U-^13^C]-glycerol are highlighted by a red arrow. **d**, Continuous growth, with daily dilutions to prevent high cell density, of the parental pleomorphic AnTat1.1E, knockout and rescue cell lines, as well as the monomorphic Lister 427 strain. The cells were grown in CMM_Glc10, CMM_Glc10/SIF, CMM_Glc0.5/Glyc10 or spent medium from AnTat1.1E culture in CMM_Glc0.5/Glyc10 maintained at low cell density to prevent SIF production (CMM_Glc0.5/Glyc10/smLCD). The skull symbolizes the death of the entire cell population. **e**, PAD1 expression levels, monitored by cell cytometry, at day three under incubation in the different growth conditions. (****, p < 0.0001; ns, not significant). **f**, Co-culture setup to determine the presence or not of a secreted (paracrine) signal involved in the production of stumpy-QS (CMM_Glc10) or stumpy-Glyc (CMM_Glc0.5/Glyc10) forms. Parental AnTat1.1E (WT) cells were consistently placed in the well as donor cells in all experiments, while the insert contained the parental AnTat1.1E (WT), △*gk* or △*aqp1-3* cell lines, which served as sentinel cells. The cultures of the donor (solid lines) and sentinel (dashed lines) cells were maintained at low cell density by daily dilution to 10^5^ cells/ml to avoid SIF production, except in the central panel where the donor cells are kept at high cell density (10^6^ cells/ml) to ensure SIF production. The diagrams on the right illustrate the co-culture setup used in each condition. Data are presented as the mean ±SD of three independent replicates. Error bars smaller than symbols are not displayed for clarity.

### The production of stumpy-Glyc forms does not require a paracrine signal

Quorum sensing is mediated by a mixture of parasite-derived extracellular di- and tripeptides (SIF), which constitutes a paracrine signal that acts on all nearby slender BSF [12,13]. To determine whether the production of stumpy-Glyc forms could be mediated by a similar paracrine signal, slender forms were cultured in spent medium from slender forms maintained at low cell density (smLCD) in CMM_Glc0.5/Glyc10 by daily dilution to avoid any SIF-induced differentiation into stumpy-QS (CMM_Glc0.5/Glyc10/smLCD in Fig.3d-e). As expected, the SIF-rich spent medium from stumpy-QS (CMM_Glc10/SIF) induced a growth arrest and PAD1 expression, whereas incubation with fresh CMM_Glc10 medium (CMM_Glc10) did not (Fig.3d-e). Spent medium from stumpy-Glyc forms maintained at low cell density (CMM_Glc0.5/Glyc10/smLCD) and fresh CMM_Glc0.5/Glyc10 medium (CMM_Glc0.5/Glyc10) induced the production of stumpy-Glyc, due to the presence of glycerol in both conditions (Fig.3d-e). To rule out the possibility of a glycerol-induced paracrine signal, the Δ*gk* and Δ*aqp1-3* mutants, which no longer respond to glycerol, were tested under the same conditions (Fig.3d-e). The growth of the Δ*gk* and Δ*aqp1-3* slender forms was not affected by the addition of CMM_Glc0.5/Glyc10/smLCD, suggesting that any putative glycerol-derived signal responsible for differentiation to stumpy-Glyc was not present in the medium. In other words, no external paracrine signal seemed to be involved in glycerol-induced differentiation (Fig.3d-e). As a negative control, the monomorphic strain *T. brucei* Lister 427, which does not respond to either SIF or glycerol, was unresponsive to the CMM_Glc10/SIF and CMM_Glc0.5/Glyc10/smLCD (Fig.3d).

To confirm these data, we performed co-culture experiments by separating donor (parental AnTat1.1E) and sentinel (parental, Δ*gk* or Δ*aqp1-3*) cell lines, using transwells. As expected, maintaining cells at low cell density in CMM_Glc10 prevented the three sentinel cell lines from differentiating into stumpy-QS (Fig.3f, left), whereas maintaining the donor cells at high density in CMM_Glc10 induced the three sentinel cells to differentiate into stumpy-QS (Fig.3f, center). At low cell density, both the parental donor and sentinel cells differentiated to stumpy-Glyc under CMM_Glc0.5/Glyc10 conditions, but the glycerol-unresponsive sentinel Δ*gk* and Δ*aqp1-3* cells still grew as slender forms under the same conditions (Fig.3f, right), confirming that no secreted paracrine signal is required for differentiation to stumpy-Glyc.

### Stumpy-Glyc cells are competent for differentiation into the procyclic insect stage

After ingestion by a tsetse fly during a blood meal, stumpy forms rapidly differentiate into the dividing procyclic insect stage, PCF, which initiates further parasite development in the insect vector. To evaluate the ability for stumpy-glyc forms to differentiate as compared to stumpy-QS forms, we first produced stumpy-glyc and stumpy-QS forms with high PAD1 expression levels by incubating slender forms in high cell density (HCD) in CMM_Glc10, low cell density (LCD) in CMM_Glc10/BHI, low cell density in CMM_Glc10/CPT, or low cell density in CMM_Glc0.5/Glyc10 for 3 days (Fig.4a-b). The cells were then transferred to 27°C in glucose-free SDM79 medium containing 6 mM cis-aconitate (CA) according to the standard differentiation procedure to PCF (Fig.4a). After 6 h of incubation in glucose-free/CA SDM79 (Fig.4a), 80% of the stumpy-QS cells as expected (*i*) became EP-positive (Fig.4c), (*ii*) exhibited a PCF morphology, *i.e.* with a longer and thinner cell body than BSF, (Fig.4d) and (*iii*) established as a stable growing population in PCF medium (Fig.4e). In comparison, CMM_Glc0.5/Glyc10-induced stumpy-glyc forms differentiated just as efficiently (Fig.4e), demonstrating that they have equivalent competency for the PCF transition. As expected, PAD1 negative slender forms maintained at low cell density (LCD) in CMM_Glc10 failed to differentiate into PCF and eventually died in glucose-free SDM79 medium (Fig.4e). As expected, the Δ*gk* slender cell line unable to differentiate into stumpy-glyc was also unable to differentiate into PCF after three days of culture in CMM_Glc0.5/Glyc10, whereas it behaved like the parental cell line when incubated at high cell density in CMM_Glc10 or at low cell density in in CMM_Glc10/SIF where it was able to differentiate into stump-QS (Fig.4f-j).

**Figure 4.**
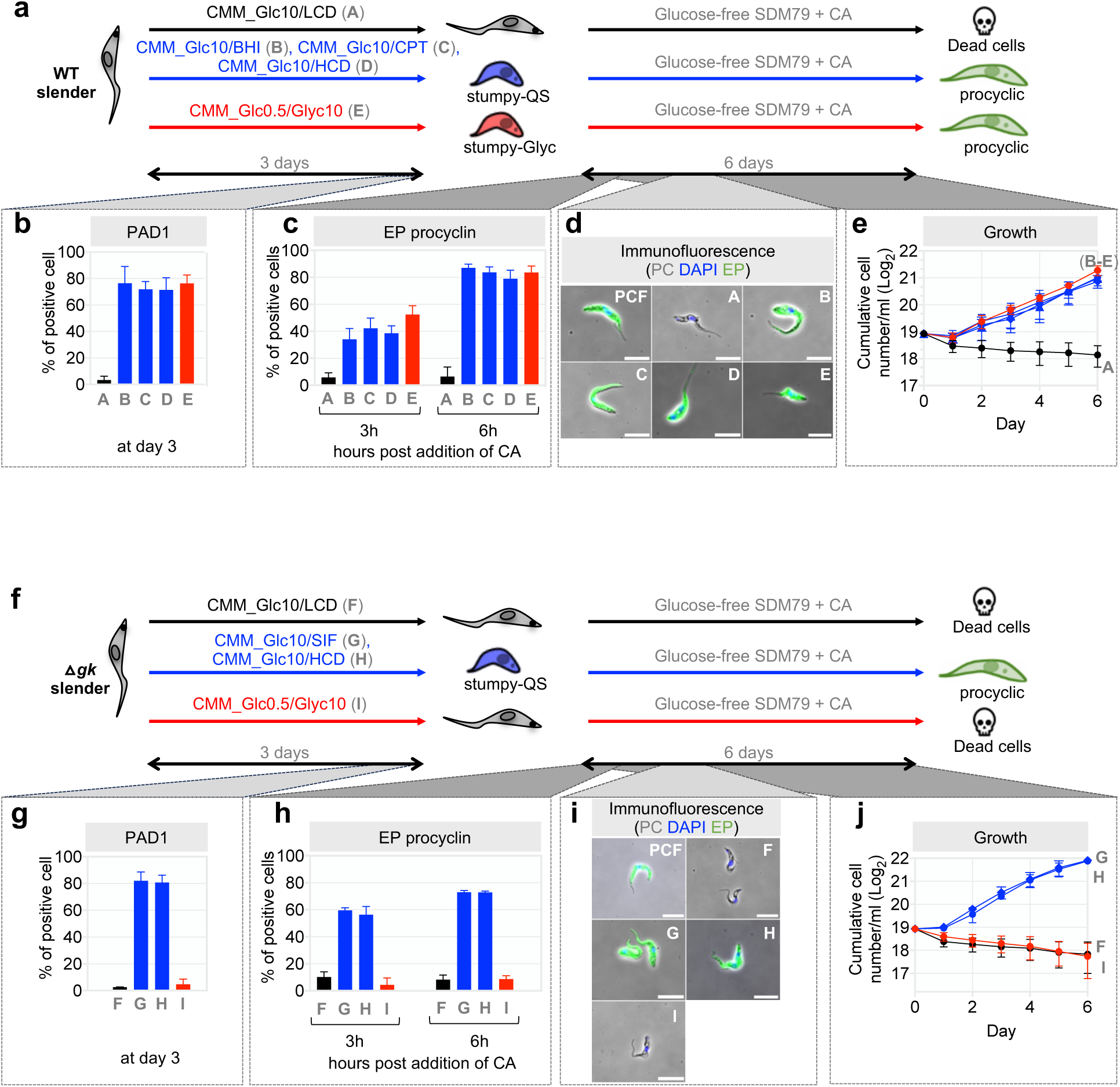
Stumpy-Glyc are competent for *in vitro* differentiation into PCF. **a,** Description of the experimental setup for the parental (WT) slender forms (AnTat1.1E-GFP::PAD1_UTR_) under conditions labelled as “A-E”. In condition “A”, the parasites were maintained as slender forms in CMM_Glc10 by daily dilution to prevent at high cell density (LCD). In conditions “B”, “C” and “D” slender forms were differentiated into stumpy-QS forms in CMM_Glc10 containing 5% of the SIF-rich BHI medium (CMM_Glc10/BHI), containing 0.1 mM 8-pCPT-cAMP (CMM_Glc10/CPT), or at high cell density without daily dilution (HCD). In condition “E” the parasites were cultivated in CMM_Glc0.5/Glyc10, for differentiation into stumpy-Glyc. This was followed by a six-day incubation in glucose-free SDM79 medium in the presence of 6 mM cis-aconitate (CA) at 27°C to induce differentiation of stumpy forms into PCF. Data are presented in panels **b-e**, *i.e.* expression of stumpy-specific marker (PAD1) by cell cytometry (**b**), expression of PCF-specific marker (EP) by cell cytometry (**c**), immunofluorescence analysis with anti-EP (**d**), and cumulative growth in the PCF glucose-free SDM79 medium over six days (**e**). **f**, Description of the experimental setup for the △*gk* mutant slender forms under conditions labelled as “F-I”. Slender forms were maintained at low cell density (“F”, CMM_Glc10 (LCD)), differentiated into stumpy-QS (“G”, CMM_Glc10/SIF; “H”, CMM_Glc10 (HCD)) or differentiated into stumpy-Glyc (“I”, CMM_Glc0.5/Glyc10) during three days, followed by a six days incubation in glucose-free SDM79 medium in the presence of 6 mM CA at 27°C to induce differentiation into PCF. Data are presented in panels **g-j**, as indicated for panels **b-e**. Data are presented as the mean ±SD of three independent replicates. Error bars smaller than symbols are not displayed for clarity.

### Glycerol is efficiently metabolized under *in vivo*-like large excess of glucose

Interstitial fluids contain ∼4 mM glucose and between 0.2 and 1 mM glycerol, depending on the tissue, while serum contains ∼10-times less glycerol (25-100 μM) [7,8]. Given that slender-to-stumpy differentiation may depend on both glycerol and glucose amounts, we investigated the effect of glycerol and glucose on the central metabolism of parasites over a range of glycerol concentrations from 0.025 to 4 mM, while maintaining a constant glucose level of 4 mM. Slender BSF were cultured with [U-^13^C]-glycerol and non-enriched glucose to enable quantification of the ^13^C-enriched and non-enriched end products excreted from glycerol and glucose metabolism (mainly pyruvate), respectively, by NMR spectrometry [22] (Fig.5a). Under Glc4/Glyc4 condition, both carbon sources contributed equally to the central carbon metabolism. It is worth noting that, under these conditions, glycerol (3 carbons) is consumed at twice the rate of glucose (6 carbons), resulting in the same amount of pyruvate being excreted. Notably, an 8-fold reduction in the glycerol amounts (Glc4/Glyc4 *versus* Glc4/Glyc0.25) mimicking tissue conditions had little effect on its metabolic flux, with only a 2-fold reduction, indicating the importance of glycerol as a primary carbon source, even in excess of glucose. However, under Glc4/Glyc0.025 conditions, representing blood conditions, the contribution of glycerol was negligible.

**Figure 5.**
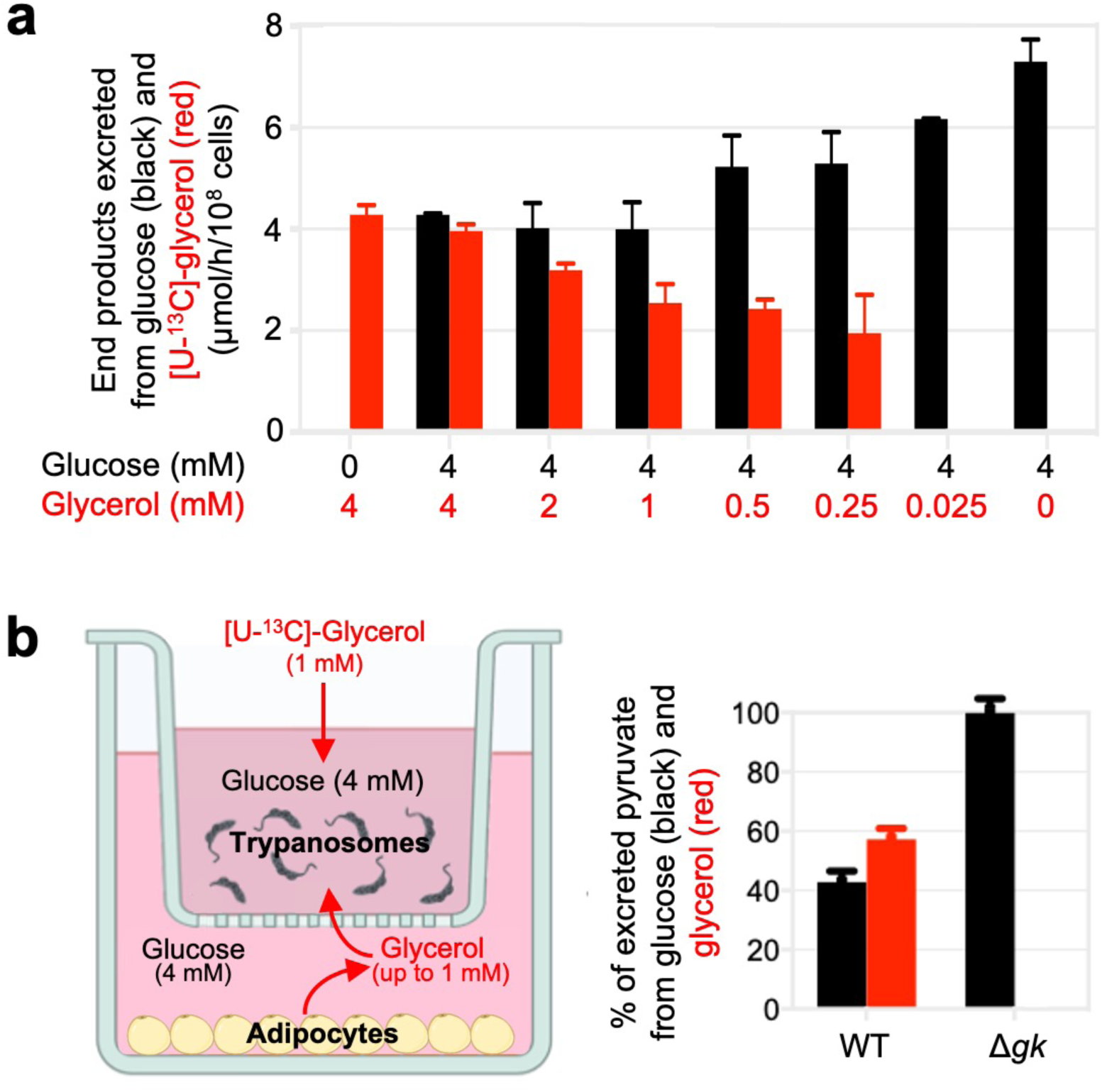
Glycerol is metabolized even in the presence of a large excess of glucose. **a,** The parental AnTat1.1E-GFP::PAD1_UTR_ slender forms were incubated 1 h in PBS containing 4 mM glucose (Glc4), 4 mM [U-^13^C]-glycerol (Glyc4) or 4 mM glucose plus 0.025, 0.25, 0.5, 1, 2 or 4 mM glycerol (Glc4/Glyc0.025 to Glc4/Glyc4). The amounts of non-enriched and ^13^C-enriched metabolic end products excreted in the spent medium from the metabolism of glucose and [U-^13^C]-glycerol, respectively, were determined by ^1^H-NMR spectroscopy. **b,** BSF trypanosomes metabolize very efficiently glycerol from adipocytes even in excess of glucose. The left panel shows an illustration of the transwell assay, which was employed in the co-culture experiment. Adipocytes are placed in the lower compartment (well), while BSF parasites are placed in the top compartment (transwell). The compartments are separated by a semi-permeable support that only enables nutrients to travel between them, preventing parasites from crossing. Uniformly ^13^C-labelled glycerol ([U-^13^C]-glycerol) was introduced at a concentration similar to that produced by adipocytes (1 mM), and glucose was adjusted to 4 mM. 24 h later, supernatants were recovered and the amounts of non-enriched and ^13^C-enriched pyruvate excreted in the spent medium from the metabolism of glucose, glycerol and [U-^13^C]-glycerol, respectively, were determined by ^1^H-NMR spectroscopy. The red circle indicates the hydrogen atoms of the methyl group that will produce peaks in the NMR spectrum. The right panel shows the contribution of either glucose or glycerol to the production of pyruvate in the two cell lines in co-culture with adipocytes (n=3 biological replicates).

This metabolic flexibility was further validated in a co-culture model using 3T3-F442a adipocytes and slender BSF (both parental and Δ*gk* cell lines) in a transwell assay. Differentiated adipocytes were first cultured alone in CMM with 20 mM glucose for 24 h to stimulate glycerol production. After this period, glucose was adjusted to 5 mM, and glycerol levels, measured enzymatically, were found to be around 1 mM. Slender BSF (parental or Δ*gk* cell lines) were then introduced into the insert, and [U-^13^C]-glycerol was added at a concentration similar to the non-enriched glycerol produced by adipocytes (Fig.5b). This setup enabled tracking of parasite-specific end-products from [U-^13^C]-glycerol metabolism. By measuring the ratio of ^12^C- and ^13^C-pyruvate relative to the total glycerol consumption, we found that in the parental cell line, glucose and glycerol contributed equally to pyruvate production, despite glucose being in excess. In contrast, in the Δ*gk* null mutant, which lacks the ability to metabolise glycerol, pyruvate was exclusively derived from glucose (Fig.5b), confirming glycerol’s significant role in the parasite’s central carbon metabolism, even in the presence of excess glucose.

### Tissue-like glycerol conditions are sufficient to trigger forms capable of infecting tsetse flies

In order to determine whether the glycerol-dependent differentiation pathway observed above could occur under physiological conditions, we examined BSF differentiation under interstitial fluid and serum concentrations of glucose and glycerol. We cultured BSF in CMM_Glc (0.5 or 4 mM glucose) containing a range of 0.01 mM to 4 mM glycerol (CMM_Glc4/Glyc0.01 to CMM_Glc4/Glyc4) (Fig.6a). At equimolar amounts of glucose and glycerol (CMM_Glc4/Glyc4, condition H), as well as under low glucose conditions (CMM_Glc0.5/Glyc4, condition I), slender BSF differentiated into growth-arrested stumpy-Glyc forms, which were mostly PAD1 positive (37% and 58%, respectively) and differentiated into PCF after transfer to 27°C in glucose-free/CA SDM79 (Fig.6b-g, H-I conditions). In contrast, in the presence of 0.01 to 0.05 mM glycerol (CMM_Glc4/Glyc0.05), slender BSF failed to differentiate into stumpy-Glyc (Fig.6b-d, B-D conditions) and then into PCF (Fig.6e-f), as shown by their slender-like growth rates and GFP::PAD1_UTR_, PAD1 and EP expression. Interestingly, an initial incubation with only 0.1 to 1 mM glycerol followed by transfer to glucose-free/CA SDM79 was sufficient to induce expression of the EP procyclin PCF marker in 20-50% of the cells, which established themselves as proliferative PCF (Fig.6e-g, E-G conditions). Very unexpectedly, under these initial conditions (0.1-1 mM glycerol), slender BSF remained highly proliferative (Fig.6b), while maintaining a slender-like morphology (Fig.S10) and showing no significant expression of the stumpy-specific marker PAD1 (Fig.6c-d), clearly showing that they are not growth-arrested stumpy-like forms. These data suggest that glycerol-sensitised slender-like forms constitute a new parasitic form virtually competent for transmission to the fly.

**Figure 6.**
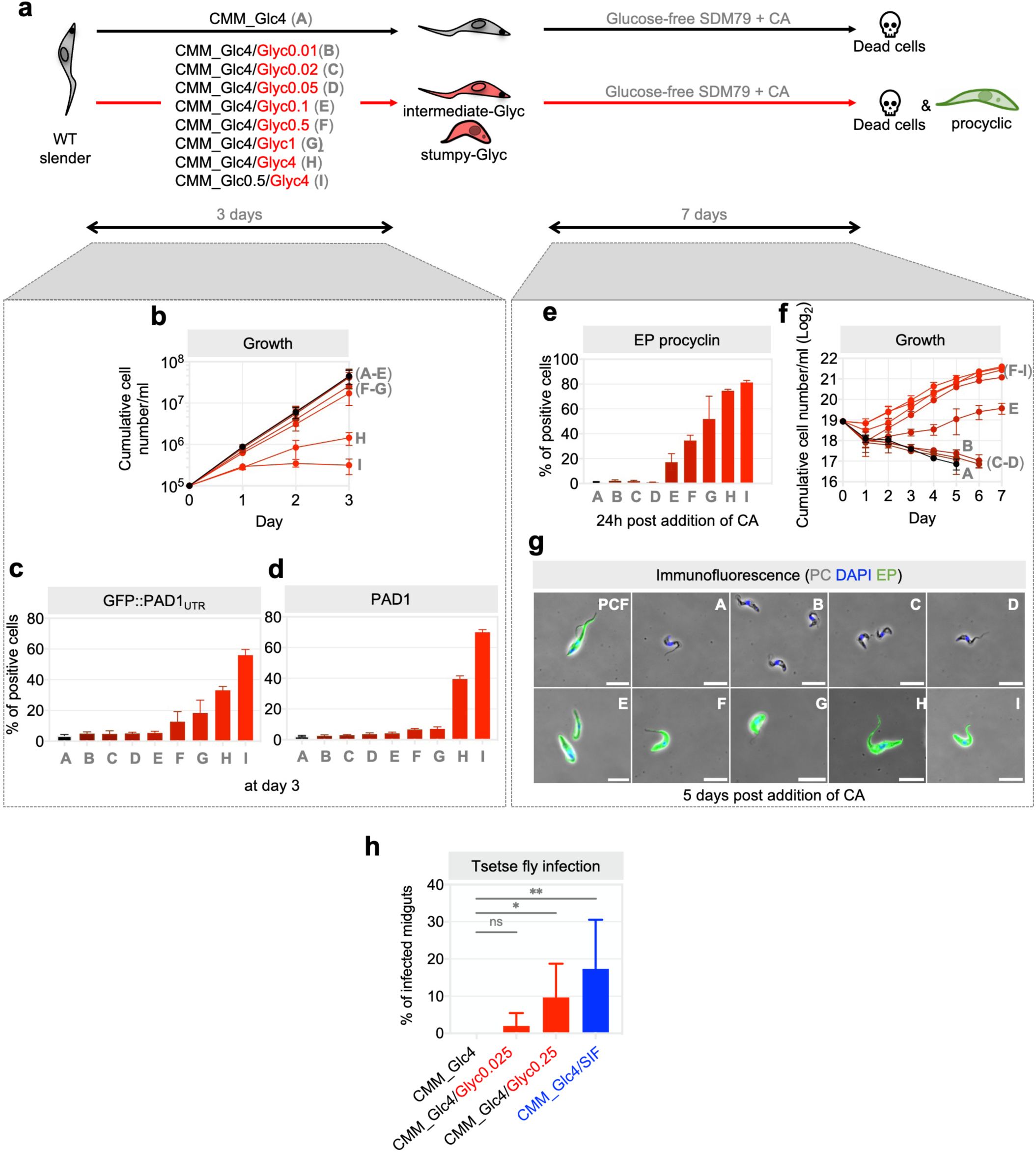
Preincubation under tissue-like conditions (low glycerol concentration) is sufficient to produce cells competent for tsetse fly infection and PCF differentiation. **a**, Description of the experimental setup for the parental (WT) slender forms (AnTat1.1E-GFP::PAD1_UTR_) under conditions labelled as “A-I”, *i.e.* CMM containing 4 mM glucose (CMM_Glc4; “A”), CMM_Glc4 containing 0.01, 0.02, 0.05, 0.1, 0.5, 1 or 4 mM glycerol (CMM_Glc4/Glyc0.01 to CMM_Glc4/Glyc4; “B” to “H”, respectively) and CMM containing 0.5 mM glucose and 4 mM glycerol (CMM_Glc0.5/Glyc4; “I”). In all these conditions, the slender forms were maintained at low cell density (LCD). After three days in CMM media, the parasites were cultivated at 27°C in glucose-free SDM79 containing 6 mM CA. Data are presented in panels **b-g**, *i.e.* continuous growth, with daily dilutions to prevent high cell density (**b**), expression of GFP::PAD1_UTR_ and PAD1 by cell cytometry analyses at day three in CMM (**c-d**), expression of PCF-specific marker (EP) by cell cytometry at day one post addition of CA (**e**), cumulative growth in the PCF glucose-free SDM79 medium over seven days (**f**), and immunofluorescence analysis with anti-EP (**g**). According to these data, only slender forms are present under conditions A-D (0 to 0.05 mM glycerol), since no PCF are generated (**f**), proliferative (**b**) intermediate forms predominate under conditions E-G (0.1 to 1 mM glycerol) and non-proliferative (**b**) stumpy-Gly are produced under conditions H-I (4 mM glycerol). Data are presented as the mean ±SD of three independent replicates. Error bars smaller than symbols are not displayed for clarity. Source data are provided as a Source Data file. **h**, Proportions of midgut infection rates in tsetse flies infected with parasites preincubated in (*i*) CMM_Glc4 as negative controls corresponding to slender forms, (*ii*) CMM_Glc4/Glyc0.025 as blood-like condition, (*iii*) CMM_Glc4/Glyc0.25 as skin-like condition, and (*iv*) CMM_Glc4/SIF as positive controls corresponding to stumpy-QS form. Each value corresponds to the mean (±SD) of 3 independent experiments (biological replicates) and a Student t-test was performed by assuming equal variances to determine the p-value (*, p < 0.1; **, p < 0.05; ns, not significant). The values are calculated from the data presented in the Source Data File

Consistent with these data, the proteomic profiles of cells grown in CMM_Glc4/Glyc2 and CMM_Glc4/Glyc0.5 were closer to those of the stumpy forms (particularly stumpy-QS obtained in CMM_Glc10/SIF) than the slender BSF, but differed from that of the stumpy-Glyc generated from the CMM_Glc0.5/Glyc10 condition (Fig.1g). The same set of proteins was regulated in both intermediate-Glyc and stumpy forms (including stumpy-Glyc), albeit at a reduced level in the former, as illustrated by the enrichment analysis of GO terms for the biological process of differentially expressed proteins (Fig.S6). Unfortunately, we did not identify any proteins specifically expressed in intermediate-Glyc forms, that could be used as specific markers, equivalent to the stumpy-specific marker PAD1.

These data highlight that, although high glucose levels may limit the ability of glycerol to generate stumpy forms, the relatively low *in vivo* levels of glycerol present in tissues are likely sufficient to generate proliferative slender-like forms suitable for fly transmission, which we have termed intermediate-Glyc. To determine whether intermediate-Glyc forms are equally capable of being transmitted by tsetse flies as stumpy-QS, teneral tsetse flies were infected with parasites pre-incubated 3 days at low cell density in either CMM_Glc4/Glyc0.025 or CMM_Glc4/Glyc0.25 conditions, corresponding to blood and skin conditions, respectively. Parasites were also incubated 3 days at low cell density in CMM_Glc4 in the presence (spent medium) or absence (fresh medium) of SIF as positive and negative controls, respectively (Fig.6h). Notably, parasites pre-incubated in the tissue-like condition (CMM_Glc4/Glyc0.25), which corresponds to intermediate-Glyc forms, infected tsetse flies at a level comparable to stumpy-QS forms pre-incubated in CMM_Glc4/SIF (9.7% versus 17.3% infection rate, respectively). In contrast, parasites pre-incubated in the blood-like condition (CMM_Glc4/Glyc0.025) infected tsetse flies in only one replicate out of three (2% infection rate) and their infectivity was (1) notably lower than that observed in intermediate-Glyc forms pre-incubated in the tissue-like condition, but (2) similar to that observed for slender forms pre-incubated under CMM_Glc4, which were unable to infect tsetse flies at all (0% infection rate). Together, these data confirm that intermediate-Glyc forms, potentially generated in physiologically relevant tissue conditions, are competent for transmission to the insect vector.

## Discussion

Throughout its life cycle, *Trypanosoma brucei* navigates diverse environments in its mammalian hosts, including the bloodstream and extravascular compartments such as the skin and adipose tissues. In the bloodstream, glucose serves as the primary carbon source, allowing the parasites to proliferate until they reach high densities, triggering a quorum-sensing mechanism that leads to the formation of cell cycle-arrested stumpy forms (stumpy-QS) considered essential for transmission to the insect vector [23]. Our study reveals that glycerol, which is actively released by adipocytes in the skin and adipose tissue [11,24], plays a novel role in parasite biology, not only in parasite metabolism, but also in their differentiation, survival and transmission. Here, we show that glycerol at high concentrations (10 mM) with accompanying low glucose levels (0.5 mM) triggers the differentiation of slender BSF into long-lived stumpy forms (termed stumpy-Glyc). These stumpy forms mimicking stumpy-QS forms but with a strikingly extended lifespan. More importantly, under tissue-like physiological conditions (4 mM glucose and 0.25 mM glycerol), we found that glycerol supports the formation of proliferative intermediate-Glyc forms that are equally transmissible to the insect vector.

Our findings could help resolve one of the long-standing conundrums in the field of *T. brucei* research, the so-called transmission paradox, in which the low parasitaemia observed in patients lies in contrast to the assumption from classical models of transmission that quorum sensing-induced stumpy forms induced by high parasite density in the bloodstream is critical for transmission [15]. In demonstrating that glycerol-induced transmissible forms are generated at low cell density compatible with low parasitaemia observed in chronic infections, we now provide a compelling possible solution to this long-standing paradox. Based on our findings, we propose an updated transmission model (Fig.7): when bloodstream parasitemia is low, with no production of stumpy-QS forms, parasite transmissibility is maintained via glycerol-induced formation of intermediate-Glyc forms in the skin. Interestingly, there is corroborating clinical evidence, in which asymptomatic carriers and chronic patient cases frequently contain cryptic *T. brucei* populations in extravascular locations, such as the basal dermis, that serve as hidden reservoirs for transmission [5,25,26]. We propose that these cryptic parasite populations could be intermediate-Glyc forms rather than the classical stumpy-QS forms. Future studies beyond the scope of this study are needed to identify a specific molecular marker and to determine the presence of these forms *in vivo*.

**Figure 7.**
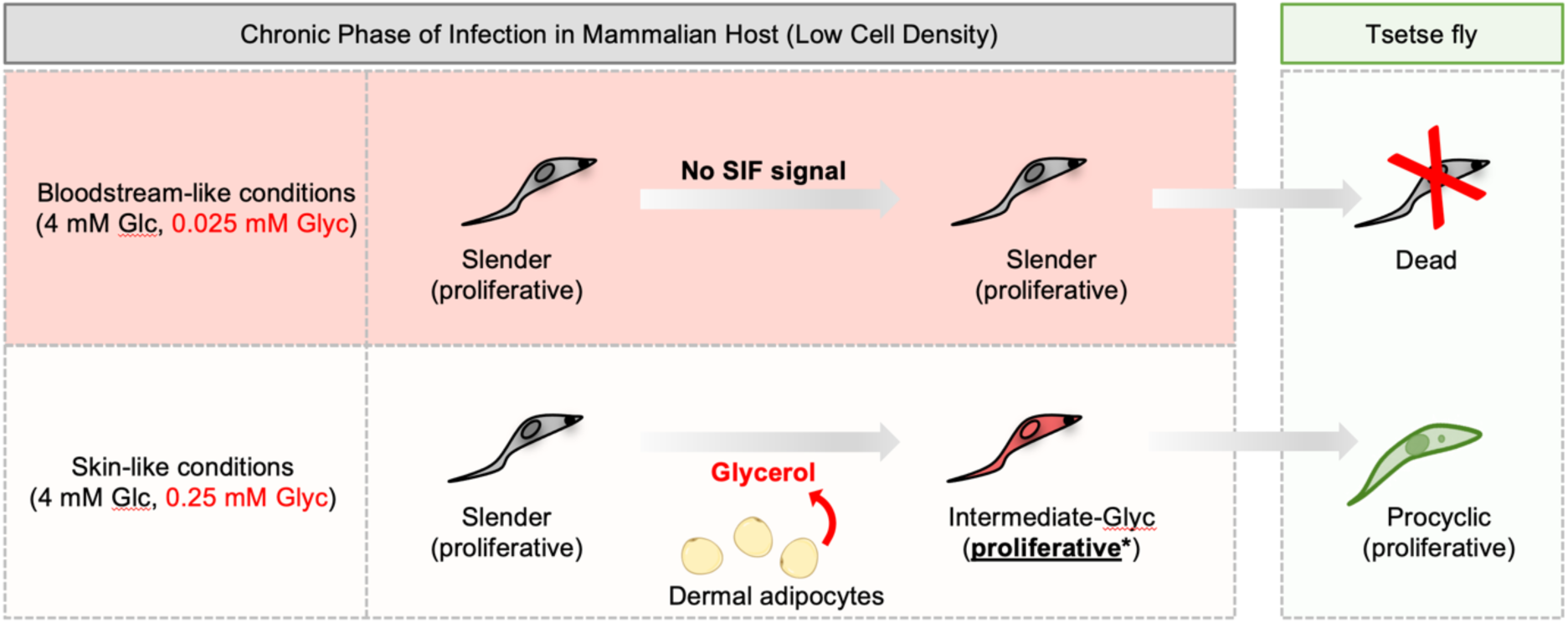
Model illustrating glycerol-driven formation of intermediate-Glyc forms in tissues and their potential transmission to tsetse flies, explaining the transmission paradox. Under low cell density conditions, typically seen during the chronic phase of infection in patients, slender forms in bloodstream-like conditions (4 mM glucose and 0.025 mM glycerol) remain proliferative, but lack ability to differentiate into procyclic forms. In this environment, these forms fail to differentiate and die when transmitted to the tsetse fly. In contrast, in skin-like conditions (4 mM glucose and 0.25 mM glycerol), slender forms encounter glycerol produced by dermal adipocytes, which induces transformation into intermediate-Glyc forms. These intermediate-Glyc forms remain <u>proliferative</u>* and display a proteome distinct from the original slender forms (Fig.1g). Importantly, they can efficiently differentiate into procyclic forms within the tsetse fly, thus providing a potential resolution to the transmission paradox.

Glucose, present at 4-5 mM in all mammalian fluids, prevents glycerol-induced differentiation into stumpy-Glyc forms from low-density slender forms and instead induce differentiation into proliferative intermediate-Glyc forms. The concomitant metabolism of glucose and glycerol probably prevents glycerol from exerting its full effect in triggering differentiation into the stumpy-Glyc forms. Under tissue-like conditions (4 mM glucose and ∼0.25 mM glycerol), glucose contributes 2.7-times more than glycerol to carbon metabolism. This moderate contribution of glycerol to metabolism is not sufficient to generate stumpy-Glyc. However, differentiation is initiated and the resulting proliferative intermediate-Glyc forms can efficiently transition into PCF *in vitro* and sustain fly infection. This contrasts with the PCF, which fully prioritize glycerol metabolism over glucose, due to the huge imbalance in activity of the glycosomal hexokinase (HK) and GK [27]. Indeed, GK activity is 74 times higher than HK activity in PCF, favoring glycerol phosphorylation over glucose phosphorylation (glycolysis), whereas this ratio is only 2 in BSF [27]. These differences in GK/HK ratio and/or the rate of glycerol uptake likely influences the differentiation process. Our proteomic analysis revealed an upregulation of aquaglyceroporins, particularly AQP1 (Supplementary Table 1), under tissue-like conditions, which may explain this dual substrate utilization. Enhanced glycerol transport via aquaglyceroporins, combined with increased glycerol kinase activity, could enable the parasite to metabolize even low levels of glycerol in excess of glucose. This might reflect an adaptive strategy that the parasite has evolved in the tissue environment to increase its capability to consume glycerol even at relatively low concentrations.

Various signals have been previously identified to trigger differentiation of stumpy forms to PCF *in vitro*, including citrate/cis-aconitate [28,29], changes in acidity or trypsin stimulation [30,31], a drop in temperature [28] and even glucose depletion [32,33]. However, these signals are ineffective in transforming proliferative slender BSF into PCF. This dogma has been recently challenged by Schuster *et al.*, who showed that proliferative slender BSF can efficiently differentiate into PCF and complete the complex life cycle in the fly, without any of the signals inducing differentiation into stumpy forms [34]. The authors proposed that their findings might help solve the paradox of parasite transmission during chronic infections. However, this idea has sparked a renewed debate in the field [23,35,36]. Based on our observations, glycerol may provide a crucial piece of evidence that could help resolve this ongoing debate. One possible explanation of the observations from Schuster *et al.*, is the high glycerol concentration (around 10 mM) found in the “trypanosome differentiation medium” commonly used for *in vitro* differentiation of BSF into PCF [37,38]. It is plausible that, when slender BSF were exposed to this medium, glycerol triggered their rapid transformation into intermediate-Glyc forms. They may have overlooked this parameter in their technical experiments as glycerol was not previously considered important in the differentiation process. Our data may both explain their observations and confirm that slender BSF cannot differentiate to PCF *in vitro*, unless triggered by glycerol or other stimuli.

In conclusion, our study highlights the complex role of glycerol in the life cycle of *Trypanosoma brucei*, revealing its multifaceted impact on parasite differentiation, survival, and transmission. Our findings not only provide a possible explanation for how *T. brucei* can persist and continue transmission even in hosts with low parasitaemia, but open the door for future work examining the skin as a reservoir for intermediate-Glyc forms that would remain available for take up by tsetse flies over days, ensuring sustained transmission.

## Acknowledgment

We thank Keith Matthews (Edinburgh, Scotland), for providing us with the anti-PAD1 immune sera. We would also like to thank Michael Boshart (Munich, Germany) for his constructive discussions on the project and the manuscript. We thank the TBMCore facility (FACSility, CNRS 3427, INSERM US005, Université de Bordeaux) for technical assistance, data acquisition on BD FACSAriaTM III Sorter, and interpretation. We also thank Marie Bao (Boston Massachusetts, USA), Scientific Editor in Life Science Editors, for science editing. FB was supported by the Centre National de la Recherche Scientifique (CNRS, https://www.cnrs.fr/), the Université de Bordeaux (https://www.u-bordeaux.fr/), the Agence National de Recherche (ANR, https://anr.fr/) ADIPOTRYP (grant number ANR-19-CE15-0004-01), TrypaDerm (grant number ANR-18-CE15-0012), and TRYPADIFF (grant number ANR-23-CE15-0040-01), and the Laboratoire d’Excellence (https://www.enseignementsup-recherche.gouv.fr/cid51355/laboratoires-d-excellence.html) through the LabEx ParaFrap (grant number ANR-11-LABX-0024) and the LabEx IBEID (grant number ANR-10-LABX-62-IBEID).

## Materials and Methods

### Trypanosomes cell cultures

The majority of the experiments described in this work were performed using the pleomorphic bloodstream form of *Trypanosoma brucei* AnTat1.1E cell line derived from a strain originally isolated from a bushbuck in Uganda in 1966 [39] and the stumpy reporter cell line GFP::PAD1_UTR_ (AnTat1.1E-GFP::PAD1_UTR_) [6,40]. The AnTat1.1E-GFP::PAD1_UTR_ cell line is derived from the AnTat1.1E 90:13 (derived from the AnTat1.1E parental cell line), a transgenic cell-line constitutively expressing the bacteriophage T7 RNA polymerase (T7RNAPol) and the Tetracycline repressor (TetR) [27], by introducing the GFP::PAD1_UTR_ DNA fragment in the tubulin intergenic region [6]. We also used the bloodstream form of *T. brucei* Lister 427 90:13, named here Lister 427, which is a monomorphic strain derived from antigenic type MiTat 1.2, clone 221a [41]. Bloodstream forms were cultured *in vitro* at 37°C in CMM (Creek’s minimal medium) supplemented with 10% (v/v) heat-inactivated fetal calf serum (FCS) [17]. The CMM was prepared as described before [17] without glucose but with the addition of all amino acids at 0.1 mM. Cell culture grade (or high purity) components were purchased from Sigma-Aldrich. One-milliliter cultures were maintained in 24 wells plates at 37°C with 5% CO_2_. Cultures were grown to a maximum density of 5 × 10^5^ cells/ml and sub-cultured by 100- or 1,000-fold dilution every 2 or 3 days, respectively. Glucose-depleted CMM (CMM_Glc0.5/Glyc10), which contains 0.5 mM coming from 10% FCS, was prepared by replacing glucose (10 mM) with 10 mM glycerol. Cell counts were obtained with a Guava EasyCyte Flow Cytometer (Merck Millipore). The AnTat1.1E-GFP::PAD1_UTR_ cell line was cultivated in CMM containing hygromycin (5 µg/ml), neomycin/G418 (2.5 µg/ml) and blasticidin (5 μg/ml) and the 427 cell line was cultivated in CMM containing hygromycin (5 µg/ml), and neomycin/G418 (2.5 µg/ml).

To obtain SIF-rich spent medium, trypanosomes were cultured in fresh CMM with 10 mM glucose at an initial density of 5 × 10⁵ cells/ml. After 48 h of growth (stationary phase), the cultures were centrifuged to pellet cells, and the supernatant was passed through a 0.22 μm filter to ensure sterility and then diluted 1:2 with fresh CMM medium. The resulting SIF-rich CMM was used for subsequent experiments.

The procyclic form of *T. brucei* EATRO1125.T7T strain [42] was cultured at 27°C in SDM79 medium containing 10% (v/v) heat inactivated fetal calf serum (FCS) and 5 μg/ml hemin [43] and in presence of hygromycin (25 µg/ml) and neomycin (10 µg/ml).

### Generation of mutant cell lines

#### 1) Glycerol kinase null mutant and rescue cell lines

Inactivation of all the *glycerol kinase* (*GK*) gene family, which is composed to 5-6 tandemly arranged almost identical copies, was performed using the CRISPR-cas9 approach as described recently [44]. Briefly, the AnTat1.1E BSF strain was transfected with a ribonucleoprotein complex composed of 30 μg of recombinant *Streptococcus pyogenes* Cas9 (Integrated DNA Technologies, IDT), 0.4 µmol of TracrRNA (IDT) and 0.4 µmol of an *in vitro*-synthesized guide RNA (TGCGCAGGTCATTCCAAACA-GGG) targeting the entire multigenic family (IDT), plus 1 µg of a repair cassette including the phleomycin resistance marker (BLE). Cells were cloned using a cell sorter (TBM Core facility) and selection of inactivated cells was performed by DNA extraction using the NucleoSpin Blood kit (Macherey-Nagel) followed by PCR amplification using primers flanking the Cas9 cleavage site. The resulting mutant cell line Δ*gk::BLE*/Δ*gk::BLE*, is named Δ*gk*.

The rescue cell line was subsequently generated by integrating into the rDNA promoter locus the pTSARib plasmid [45] containing the particular *GK* gene, fused at the N-terminal end with BLA as a resistance cassette. Rescue BSF trypanosomes were generated using a Nucleofector (Lonza) as described previously [46,47]. Selection-marker recovery was confirmed by screening individual clones in multi-well plates. Transformants were selected with the appropriate antibiotic concentrations: phleomycin (2.5 μg/ml) and blasticidin (5 μg/ml). Clonal populations were obtained by limiting dilution and cell culture growth was monitored with an automatic Guava EasyCyte Flow Cytometer (Merck Millipore). The resulting Δ*gk::BLE*/Δ*gk::BLE*/*GK::BLA* mutant cell line, is named Δ*gk*/GK.

#### 2) Aquaglyceroporin null mutant and rescue cell lines

For generating the *AQP1-2-3* null knockout, and the add-back cell line, all insert templates were synthesized by GeneCust Europe (Dudelange, Luxembourg). Aquaglyceroporin-flanking sequences were inserted on both sides of selectable marker cassettes. Both alleles of the *AQP1* gene (966 bp, AJ697889, Tb427.06.1520) were replaced by NEO and BLE resistance cassettes for the engineering of parasites lacking *AQP1*. For deleting *AQP2* (AJ697890, Tb427.10.14170) and *AQP3* (AJ697891, Tb427.10.14160) genes which are arranged as a tandem pair on chromosome 10, both alleles of the *AQP2-3* locus were entirely disrupted by replacing the 6,363 bp fragment with HYG and PAC markers. For creating the *AQP1-2-3* triple null trypanosomes, the *AQP2-3* locus was successively disrupted in *AQP1* KO parasites, yielding parasites resistant to four resistance cassettes (NEO, BLE, HYG and PAC).

The add-back cell line was subsequently generated by integrating into the rDNA promoter locus the pTSARib plasmid [48] containing the particular *AQP* gene(s) fused to the EGFP fluorescent marker at the N-terminal end, and BLA as a resistance cassette. Specifically, the following rescue cell line was generated: *AQP1-2-3* null parasites expressing *EGFP-AQP1*.

PCR amplifications of the DNA fragments bearing the distinct *AQPs* flanking sequences and the appropriate resistance markers were used for nucleofection and generation of *AQP1-2-3* null cell line. The primers used are listed below: 5’-AAGAAAGAAAATTATTCAGATATA-3’ (FW, located 24 bp upstream the flanking 5’UTR sequence of *AQP1* used for the first allele replacement by NEO); 5’-GGAGAGGAAGGGGAGAAAAGGGAG-3’ (RV, last 24 bp downstream the flanking 3’UTR sequence of *AQP1* used for the second allele replacement by BLE); 5’-TTTCACTCCTCTACGCACACGC-3’ (FW, located 22 bp upstream the flanking 5’UTR sequence of *AQP2* used for the first allele replacement by HYG); 5’-TTCACCTTATTTATCTCTAAAAATTAG-3’ (RV, last 27 bp downstream the flanking 3’UTR sequence of *AQP3* used for the second allele replacement by PAC). On the other hand, add-back plasmids were linearized with *Sph*I prior to nucleofection. The resulting Δ*aqp1::NEO*/Δ*aqp1::BLE*/Δ*aqp2-3::HYG*/Δ*aqp2-3::PAC* and Δ*aqp1::NEO*/Δ*aqp1::BLE*/Δ*aqp2-3::HYG*/Δ*aqp2-3::PAC*/EGFP-*AQP1::BLA* mutant cell lines, are named Δ*agp1-3* and Δ*agp1-3*/AQP1, respectively.

Knockout and rescue BSF trypanosomes were generated using a Nucleofector (Lonza) as described previously [46,47]. Selection-marker recovery was confirmed by screening individual clones in multi-well plates. Transformants were selected with the appropriate antibiotic concentrations: neomycin/G418 (1.5 μg/ml), phleomycin (2.5 μg/ml), hygromycin (5 μg/ml), puromycin (0.1 μg/ml), blasticidin (5 μg/ml) and nourseothricin (25 μg/ml). Clonal populations were obtained by limiting dilution and cell culture growth was monitored with an automatic Guava EasyCyte Flow Cytometer (Merck Millipore).

### RT-qPCR validations

#### i) RNA isolation

Total RNA from samples was isolated using the Qiagen RNeasy Mini Kit (Qiagen). For each cell line, a total of 2.5 × 10^7^ parasites from culture were centrifuged at 1,400 RCF and washed twice with TDB (Trypanosome Dilution Buffer). The pellets were immediately resuspended in 350 μl of lysis buffer (RLT buffer + Beta-mercaptoethanol) and samples were homogenized by pipetting until complete resuspension. The rest of the extraction was proceeded according to the manufacturer’s instructions. The concentration of each RNA sample was measured by spectrophotometric analysis in a NanoDrop 2000c (ThermoFisher Scientific). Finally, RNA quality was determined by capillary electrophoresis in a 2100 Bioanalyzer (Agilent). Extracted RNA was stored at −80 °C prior to RT-qPCR analyses.

#### ii) DNAse treatment and validation

Extracted RNA samples were subjected to a DNase treatment using Invitrogen’s DNA-free kit (Life Technologies) according to manufacturer’s protocol. DNase treatment confirmation was performed by running a qPCR targeting the tubulin locus as housekeeping gene control. The primer pair used is as follows: forward (FW), 5’-ACTGGGCAAAGGGCCACTAC-3’; reverse (RV), 5’-CTCCTTGCAGCACACATCGA-3’, with an amplicon size of 105 bp. Reactions were done in a volume of 20 µl containing: 1 µl of RNA template (20 ng), 10 µl of 2X GoTaq qPCR Master Mix Buffer, 2 µl of both FW and RV primers (at 10X) and 5 µl of Nuclease-Free Water. Amplification was accomplished in a QuantStudioTM 3 thermal cycler (Applied Biosystem) using the following program: 2 min at 95°C; 40 cycles at 94°C 15 sec; 55°C 1 min; and final 30 sec at 60°C. The absence of residual DNA in the DNase-treated RNA samples was thus confirmed when no amplification was observed. RNA samples were further employed for gene expression quantification.

#### iii) Primer design of target genes

Tubulin was used here as housekeeping gene, the primer sequences were described elsewhere [48] and are listed below: FW, 5’-ACTGGGCAAAGGGCCACTAC-3’; RV, 5’-CTCCTTGCAGCACACATCGA-3’. For Aquaglyceroporin 1 (*AQP1*), primer sequences were obtained using the primer design tool from NCBI (https://www.ncbi.nlm.nih.gov/tools/primer-blast/) and tested both in silico using AmplifX2 V2.1.1 (https://inp.univ-amu.fr/en/amplifx-manage-test-and-design-your-primers-for-pcr), and in vitro through both serial dilutions and gradient temperatures to respectively determine the optimal concentration and temperature. The primers are listed below: FW, 5’-ATGCGATTAATCCGGCTCGT-3’; RV, 5’-GGCCAAAAATGCCTCCCAAG-3’. Since the rescues were generated by adding-back the gene fused to the eGFP fluorescent marker at the N-terminal end, the quantification of the expression of the gene of interest in the add-back was performed by targeting the level of expression of eGFP. The primer sequences were described elsewhere (https://signagen.com/blog/2020/09/01/egfp-primers-and-probe-for-qpcr/) and are listed below: FW, 5’-AGTCCGCCCTGAGCAAAGA-3’; RV, 5’-TCCAGCAGGACCATGTGATC-3’. All primers were used at a final concentration of 250nM.

#### iv) Real time RT-PCR assay

One step RT-PCR kit (Promega) was used. Reactions were prepared in a volume of 20 µl, containing: 1 µl of RNA template (20 ng), 10 µl of 2X GoTaq qPCR Master Mix Buffer, 0.4 µl of 50X GoScriptTM RT Mix for 1-Step RT-qPCR, 2 µl of both FW and RV primers (10X), and 4.6 µl of Nuclease-Free Water. Reverse transcription and amplification were accomplished in one step in a QuantStudioTM 3 thermal cycler (Applied Biosystem) using the following incubation program: 15 min at 42°C; 10 min at 95°C; 40 cycles of 95°C during 30 sec; 55,5°C for tubulin, and 57°C for both AQP1 and eGFP, during 1 min; final 72°C during 30 sec. A melt curve program was included: 15 sec at 95°C; 55,5°C for tubulin, and 57°C for both AQP1 and eGFP during 1 min; 95°C for 10 sec. Amplicons were then analyzed by gel electrophoresis. Negative controls consisted of: (1) Reaction mix with no reverse-transcriptase + total RNA (DNase treated) extracted from wild-type trypanosomes in culture; (2), Reaction mix with no primers + total RNA (DNase treated) extracted from wild-type trypanosomes in culture; (3) Reaction mix with No template. While positive control consisted of Reaction mix with total RNA (DNase treated) extracted from wild-type trypanosomes in culture.

#### v) RT-qPCR data analyses

All samples were amplified in triplicates and Cq mean values were calculated. The level of expression of each gene were calculated according to the 2^−ΔΔCq^ algorithm.

### Anti-PAD1 immunopurified serum production

The anti-PAD1 immune serum was produced by Covalab in rabbits through four injections, administered at 20-day intervals, using specific peptides for PAD1 (H_2_N-CPKEPTRDAREAAPQ-COOH and H_2_N-ETCCRREVAE-COOH) [16] and emulsified with complete (for the first injection) or incomplete Freund’s adjuvant. After 74 days of immunization, the rabbit was sacrificed, and the immune serum collected was tested for specificity and exhibited both specific and non-specific bands. Consequently, the serum was immunopurified on each peptide separately. The immunopurification protocol involved several steps: 1 ml of activated Sepharose beads was coupled with the peptide, and 5 ml of the antiserum was incubated with these beads for 1 h at 37°C. After incubation, the column was washed multiple times, and antibodies were eluted using a buffer containing 0.02% sodium azide. The antibody concentration in the collected fractions was determined by measuring absorbance at 280 nm. ELISA testing was performed with peptides adsorbed on a microtiter plate, and antibody dilutions ranged from 1,000 ng/ml to 8 ng/ml. The titer was established as the lowest antibody concentration at which absorbance at 450 nm was ≥ 1. Antibodies were deemed specific and immunoreactive if the titer was ≤ 100 ng/ml. The final antibody concentration was 74 µg/ml, with a total purified volume of 4.2 ml. After evaluating the specificity of the two immunopurified sera, the serum purified on peptide 1 (H_2_N-CPKEPTRDAREAAPQ-COOH) was identified as specific and was thus selected for further experiments.

### Western blot analyses

Total protein extracts of parental or mutant *T*. *brucei* BSF cell lines (5 × 10^6^ cells per lane) harvested at a density of 5 × 10^5^ cells/ml (or more depending on the condition) were separated by SDS-PAGE (10%) and immunoblotted on TransBlot Turbo Midi-size PVDF Membranes (Bio-Rad) [49]. Immunodetection was performed as described [49,50], using the primary antibodies, the rabbit anti-GK (1:5,000, gift from P.A.M. Michels, Edinburgh, UK), the rabbit anti-PAD1 (an antipeptide antibody recognizing TbPAD1 amino acids CPKEPTRDAREAAPQ was generated in rabbits, by Covalab, as described before [16], and diluted 1:1,000), the rabbit anti-ASCT (1:100) [51], the rabbit anti-IVDH (1:100) [52], the mouse monoclonal anti-TAO (7D3, 1:100, gift from M. Chaudhuri, Nashville, TN, USA) [53] and the mouse anti-tubulin (1:3,000, gift from T. Noel, Bordeaux, France). Anti-rabbit or anti-mouse antibodies conjugated to the horseradish peroxidase (Bio-Rad) were used as secondary antibodies. Revelation was performed using the Clarity Western ECL Substrate as described by the manufacturer (Bio-Rad). Images were acquired and analyzed with the ImageQuant LAS 4000 luminescent image analyzer (GE Healthcare Life Sciences).

### Immunofluorescence and cell cycle analyses

Parasites grown in culture were collected by centrifugation, washed and fixed in paraformaldehyde as described elsewhere. Slides were incubated with primary antibodies followed by anti-mouse Alexa Fluor 488 secondary antibody or anti-rabbit Alexa Fluor 647 secondary antibody (diluted 1:400) (ThermoFisher). The nuclei were stained with DAPI (10 µg/ml) and cells were observed using a Zeiss Axio imager Z1 microscope; images were captured using Metamorph software (Molecular Devices). Images were processed using ImageJ software [54]. Parasite morphometric measurements were scored from phase contrast microscopy images, analyzed via the automated tool HTIAoT (http://users.ox.ac.uk/~path0493/htiaot). This automated tool processes phase contrast images to identify cells, applying a rolling ball background subtraction filter and thresholding to generate cell outlines. It analyses color-deconvolved images to measure total kinetoplast and nuclear DNA content, create organelle masks, and count the number of kinetoplasts and nuclei per cell. The Medial Axis Transform (MAT) was used to assess cell morphology, including length, maximum width, and profile shape, and to determine the position of organelles within the cell [55].

### Flow cytometry

2-5 × 10^6^ cells were washed twice in PBS prior to fixing in 500 µl 4% paraformaldehyde in PBS for 20 min at RT. Cells were then washed 3x in PBS and resuspended in 4% BSA:PBS for 30 min. Cells were then resuspended in primary antibody diluted in 4% BSA:PBS, anti-EP procyclin (Cedar Lane laboratories, diluted 1:500) or anti-PAD1 (diluted 1:500), and were incubated overnight at 4°C. The cells were washed twice in PBS and were resuspended in secondary antibody diluted in 4% BSA:PBS (anti-mouse Alexa Flour 647&488 were diluted 1:500). Cells were washed twice in PBS and were resuspended in 500 µl PBS. Samples were then processed on a Guava EasyCyte Flow Cytometer (Merck Millipore). Positive controls and secondary antibody only controls were included.

### *In vitro* differentiation to procyclic forms (PCF)

To induce the differentiation of *T*. *brucei* parasites into PCF *in vitro*, a specialized glucose-free medium (SDM79-GlcFree) was developed as described before [56]. The SDM79-GlcFree medium was formulated by excluding glucose from SDM79 (CliniSciences) through a sequential process. This medium was prepared by growing parental PCF trypanosomes (5 × 10^6^ cells/ml) in glucose-free SDM79 supplemented with 20% FCS, during 3 days to late log phase (2 × 10^7^ cells/ml), then the spent medium was filtered and completed with one volume of fresh glucose-free and FCS-free SDM79.

For the differentiation process, parasites were resuspended at a concentration of 5 × 10^5^ cells/ml in the SDM79-GlcFree medium, supplemented with 6 mM cis-aconitate (Sigma, A3412), and maintained at 27°C. The differentiation progression was assessed at distinct time points, specifically at 0, 3, 6 and 24 h, by employing flow cytometry. A pivotal indicator of the transition to PCF was the expression of EP procyclin, a surface protein characteristic of PCF. The quantification of EP procyclin-expressing parasites was performed using the previously mentioned flow cytometry method.

### DNA synthesis analyses

DNA synthesis was measured by incorporation of 5-ethynyl-2ʹ-deoxyuridine (EdU), which was subsequently detected using fluorescent azide via click chemistry [40]. Parasites pre-incubated under different conditions were treated with EdU (100 μM) for 30 min. The subsequent protocol closely followed the manufacturer’s instructions with minor modifications (Merck - Sigma Aldrich). First, parasites were washed with 1x TDB, fixed with 1% paraformaldehyde for 10 min and then quenched with 0.125 M glycine for 5 min. Parasites were then attached to coverslips for 30 min, permeabilized with ice-cold methanol for 20 min and washed twice with 4% BSA in 1x PBS. Parasites were then fluorescently labelled with Alexa Fluor® 488 fluorescent azide by incubation with 300 μl Click-iT® reaction cocktail for a further 30 min in the dark. Finally, the cells were washed twice with 4% BSA in 1x PBS, stained with 1 μg/ml DAPI for 5 min, and the percentage of EdU-positive cells was assessed by fluorescence microscopy. All steps were performed at room temperature and more than 300 cells were analyzed per condition.

### Analyses of excreted end products by proton NMR

BSF trypanosomes (10^7^ cells, ∼0.1 mg of protein) were harvested at a density of 5 × 10^5^ cells/ml by centrifugation at 1,400 g for 10 min, washed once with phosphate-buffered saline (PBS) containing 1 mM of the carbon source and incubated for 1.5 h at 37°C in 1 ml of incubation buffer (PBS supplemented with 5 g/l NaHCO_3_, pH 7.4), with [U-^13^C]-glucose or glucose (4 mM) in the presence or the absence of glycerol (4 - 0.5 mM). This quantitative ^1^H-NMR approach was previously developed to distinguish between [13C]-enriched and non-enriched excreted molecules produced from [13C]-enriched and non-enriched carbon sources, respectively [22,56]. The integrity of the cells during the incubation was checked by microscopic observation. 50 μl of maleate (10 mM) were added as internal reference to a 500 μl aliquot of the collected supernatant and proton NMR (^1^H-NMR) spectra were performed at 500.19 MHz on a Bruker Avance III 500 HD spectrometer equipped with a 5 mm cryoprobe Prodigy. Measurements were recorded at 25°C. Acquisition conditions were as follows: 90° flip angle, 5,000 Hz spectral width, 32 K memory size, and 9.3 sec total recycle time. Measurements were performed with 64 scans for a total time close to 10 min 30 sec. Resonances of the obtained spectra were integrated and metabolites concentrations were calculated using the ERETIC2 NMR quantification Bruker program.

### Label-free quantitative proteomics

Three independent biological replicates were performed on total protein extracts from different forms of trypanosomes. Proteins were extracted with 4% SDC in 100 mM Tris HCL solution, in addition of protease inhibitors, boiled 5 min and finally sonicated. The steps of sample preparation and protein digestion by the trypsin were performed as previously described [37]. NanoLC-MS/MS analysis were performed using an Ultimate 3000 RSLC Nano-UPHLC system (Thermo Scientific, USA) coupled to a nanospray Orbitrap Fusion™ Lumos™ Tribrid™ Mass Spectrometer (Thermo Fisher Scientific, California, USA). Each peptide extracts were loaded on a 300 µm ID x 5 mm PepMap C18 precolumn (Thermo Scientific, USA) at a flow rate of 10 µl/min. After a 3 min desalting step, peptides were separated on a 50 cm EasySpray column (75 µm ID, 2 µm C18 beads, 100 Å pore size, ES903, Thermo Fisher Scientific) with a 4-40% linear gradient of solvent B (0.1% formic acid in 80% ACN) in 115 min. The separation flow rate was set at 300 nl/min. The mass spectrometer operated in positive ion mode at a 1.9 kV needle voltage. Data was acquired using Xcalibur 4.4 software in a data-dependent mode. MS scans (m/z 375-1500) were recorded at a resolution of R = 120000 (@ m/z 200) and a standard AGC target, ions collected within 50 ms, followed by a top speed duty cycle of up to 3 sec for MS/MS acquisition. Precursor ions (2 to 7 charge states) were isolated in the quadrupole with a mass window of 1.6 Th and fragmented with HCD@30% normalized collision energy. MS/MS data was acquired in the Orbitrap cell with a resolution of R=30000 (@m/z 200), a standard AGC target and a maximum injection time in automatic mode. Selected precursors were excluded for 60 sec. Protein identification and label-free quantification (LFQ) were done in Proteome Discoverer 3.0. The CHIMERYS node, using the prediction model inferys_2.1 fragmentations, was used to identify proteins in batch mode by searching against a *Trypanosoma brucei* TREU 927 protein database (9243 entries, version 65). Two missed enzyme cleavages were allowed for trypsin. Peptide lengths of 7–30 amino acids, a maximum of 3 modifications, charges of 2–4, and 20 ppm for fragment mass tolerance were set. Oxidation (M) and carbamidomethyl (C) were respectively searched as dynamic and static modifications by the CHIMERYS software. Peptide validation was performed using the Percolator algorithm and only “high confidence” peptides were retained corresponding to a 1% false discovery rate at the peptide level. Minora feature detector node (LFQ) was used along with the feature mapper and precursor ions quantifier. The normalization parameters were selected as follows: (1) Unique peptides, (2) Precursor abundance based on intensity, (3) Normalization mode: total peptide amount, (4) Protein abundance calculation: summed abundances, (5) Protein ratio calculation: pairwise ratio based and (6) Missing values were replaced with random values sampled from the lower 5% of detected values. Quantitative data were considered for master proteins, quantified by a minimum of 2 unique peptides, a fold changes above 2 and a statistical p-value adjusted using Benjamini-Hochberg correction for the FDR lower than 0.05. The mass spectrometry proteomics data have been deposited to the ProteomeXchange Consortium via the PRIDE [57] partner repository with the dataset identifier PXD056405.

### Trypanosomes and adipocytes co-culture

The 3T3F442a pre-adipocyte cells (Kerafast, USA) were cultured under standard conditions using a 37°C incubator with 10% CO₂ [58]. The culture medium consisted of DMEM High Glucose (Gibco, Fisher cat#11574486) supplemented with 10% iron-supplemented bovine calf serum (GE Healthcare, Fisher cat#11551831) and 1% penicillin/streptomycin. Cells were plated at a density of 2 × 10³ to 2 × 10⁴ cells/cm² depending on their viability and flask size (T75 or T150), and were incubated overnight. The medium was changed the next day or every other day thereafter, ensuring optimal growth conditions.

For co-culture experiments, 3T3F442a pre-adipocytes were plated at 1 × 10⁴ cells/cm² in a 6-well plate (Falcon, UGAP CAT# 2515137). Once the cells reached confluence, typically after 48 h, the media was carefully removed, and 2 ml of pre-warmed DMEM supplemented with 10% fetal bovine serum (FBS) and 1% penicillin/streptomycin was added to each well. This process initiates differentiation. Media was replaced with ½ IMDM mixed with ½ DMEM, supplemented with 10% FBS, 1% penicillin/streptomycin, and 5 µg/ml insulin (Sigma, I9278). This media change continued every 48 h for 1 to 2 weeks, depending on the efficiency of differentiation.

At the start of the experimental week, approximately after 18-20 days, the media was replaced with CREEKS Minimal Media (CMM with the desired glucose concentration, 10% FBS, and 1% penicillin/streptomycin) [17]. The next day, a transwell insert (Falcon Ø 234 mm, Pores Ø 0.4 μm, UGAP CAT# 2515125) was added to introduce BSF trypanosomes (5 × 10^5^ cells/ml) into the co-culture, thereby initiating the co-culture phase of the experiment. Media aliquots were taken every day for proton NMR analysis.

### Tsetse fly maintenance, infection and dissection

*Glossina morsitans morsitans* tsetse flies of the Paris colony were maintained in Roubaud cages at 24-25°C and 60-70% humidity and fed through a silicone membrane with fresh mechanically defibrinated sheep blood (BCL, France). Fly infections and analyses have been performed as described before [23]. Briefly, teneral males (between 24 h and 72 h post-emergence) were allowed to ingest parasites through a silicone membrane and without any chemical immunomodulatory supplement. A total of 3 independent biological replicates per condition were performed with batches of 50 flies per condition. Flies were starved for at least 24 h before being dissected blindly 14 days post-ingestion for detection of PCF in the midgut.

### Statistical analyses

Statistical analysis was performed using GraphPad Prism (version 10.0.1). Data are presented as mean ± standard deviation (SD). Statistical differences were assessed using two-way analysis of variance (ANOVA) and one-way ANOVA with Šidák’s test for multiple comparisons. Stand-alone comparisons were performed using unpaired t-tests. P values lower than 0.05 were considered to be statistically significant. However, for specific experiments, a threshold of P < 0.1 was considered statistically significant, as indicated in the figure legends.

## Supplementary figures

**Figure S1.**
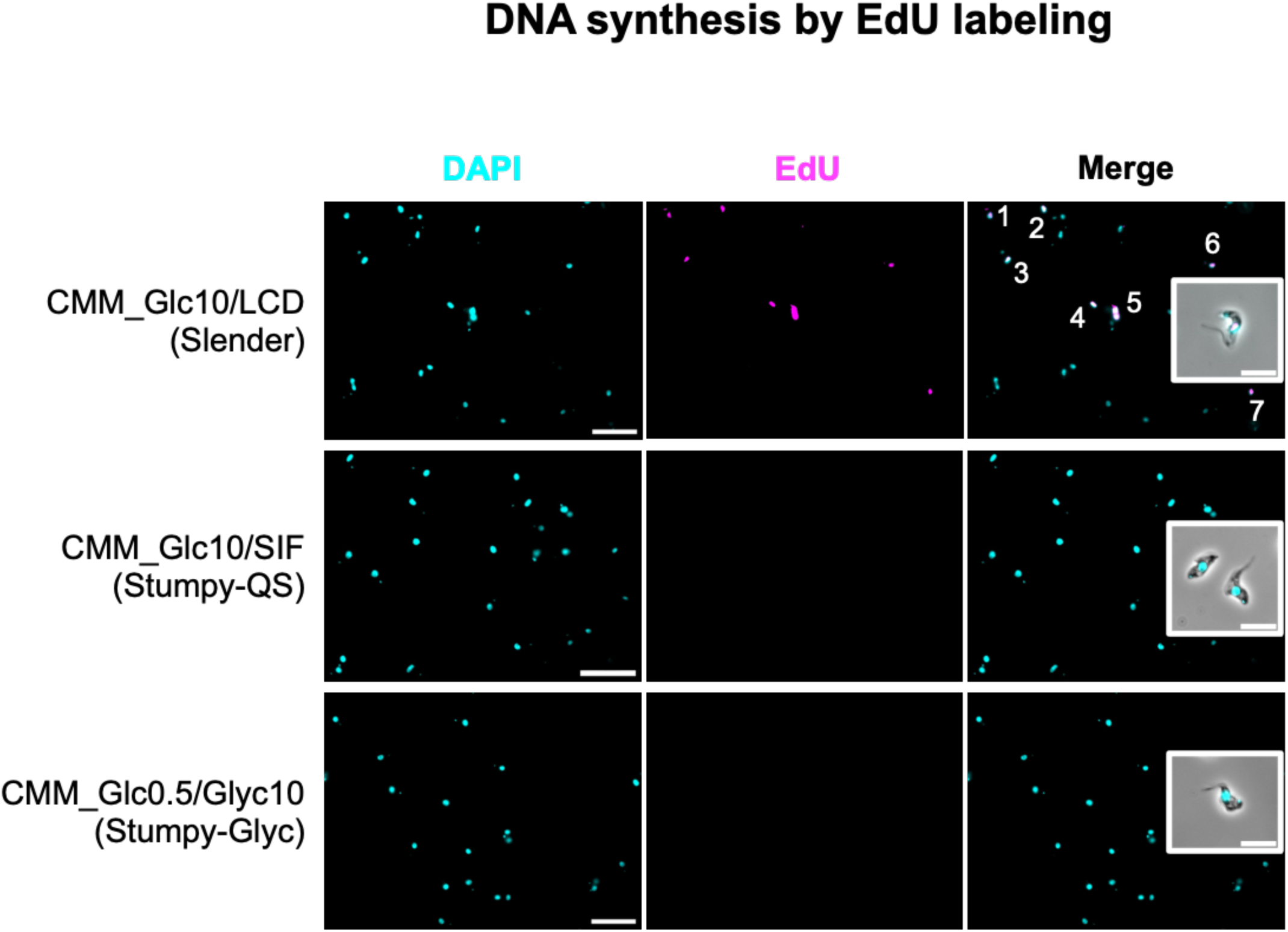
Stumpy-Glyc cells are cell-cycle arrested. Microscopy images of the AnTat1.1E-GFP::PAD1_UTR_ incubated in (*i*) CMM_Glc10 and maintained as slender forms by daily dilution to prevent high cell density (LCD), (*ii*) in SIF-rich CMM_Glc10 spent medium derived from a stationary phase culture of AnTat1.1E-GFP::PAD1_UTR_ (CMM_Glc10/SIF) and (*iii*) in low glucose (0.5 mM) CMM containing 10 mM glycerol (CMM_Glc0.5/Glyc10), with EdU (false colored magenta) and DAPI (false colored cyan) (n = 3 independent experiments). Scale bar = 20 μm. Insets: representative images of labelled parasites (scale bar, 10 μm).

**Figure S2.**
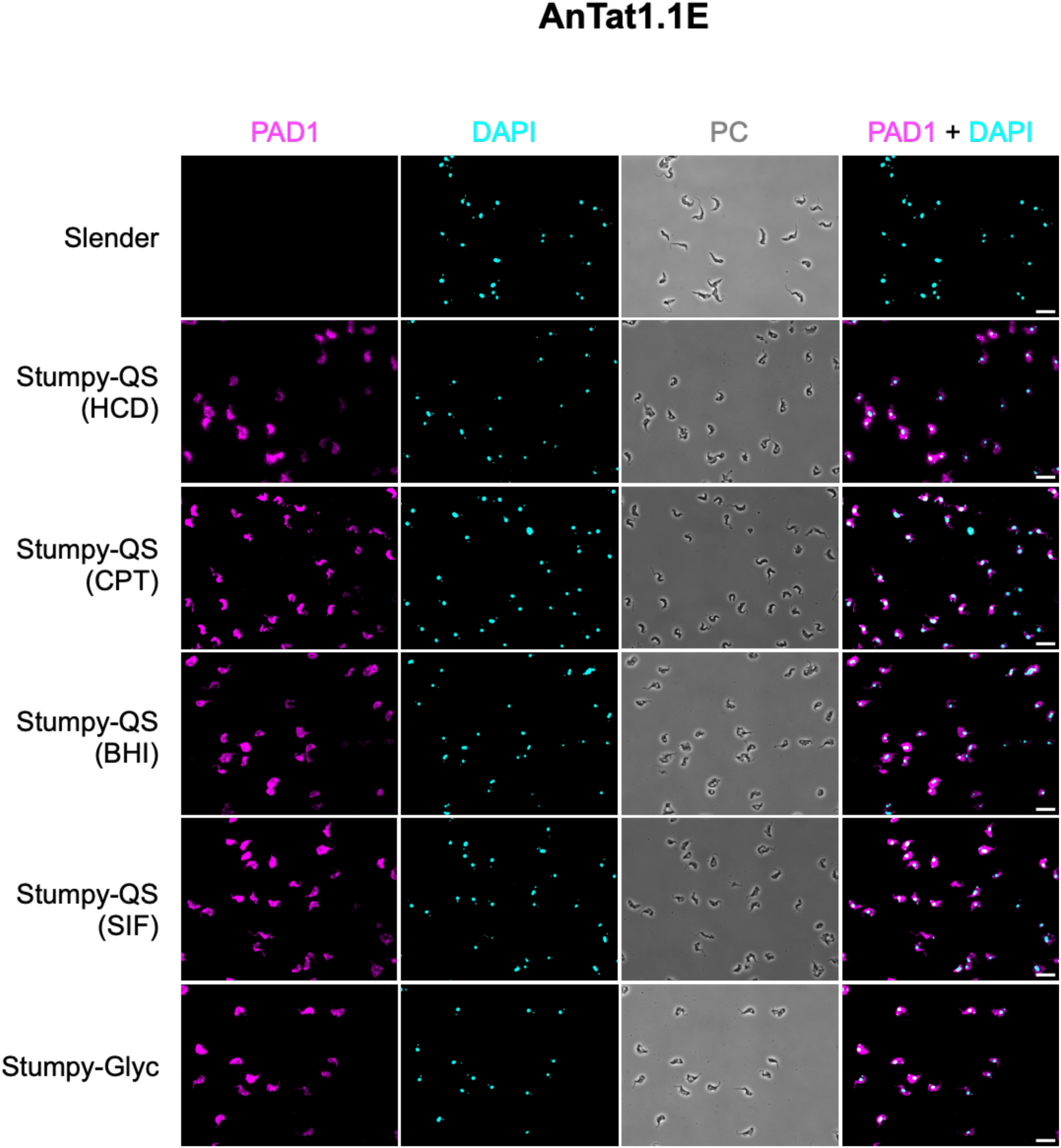
Stumpy-Glyc express PAD1 to the plasma membrane. Immunofluorescence analysis of the wild type slender cells (AnTat1.1E) maintained at low cell density (*i*), stumpy-QS generated either at high cell density in CMM_Glc10 (HCD) (*ii*), in CMM_Glc10 containing 100 µM 8-pCPT-cAMP (CPT) (*iii*), in CMM_Glc10 containing 5% BHI medium (BHI) (*iv*) and in SIF-rich CMM_Glc10 spent medium derived from a stationary phase culture of AnTat1.1E-GFP::PAD1_UTR_ (SIF) (*v*), as well as stumpy-Glyc generated in low glucose (0.5 mM) CMM containing 10 mM glycerol (*vi*). PAD1 expression is revealed using anti-PAD1 (false colored magenta), nucleus and kinetoplast using DAPI (false colored Cyan). PC Phase contrast. Scale bar = 20 µm.

**Figure S3.**
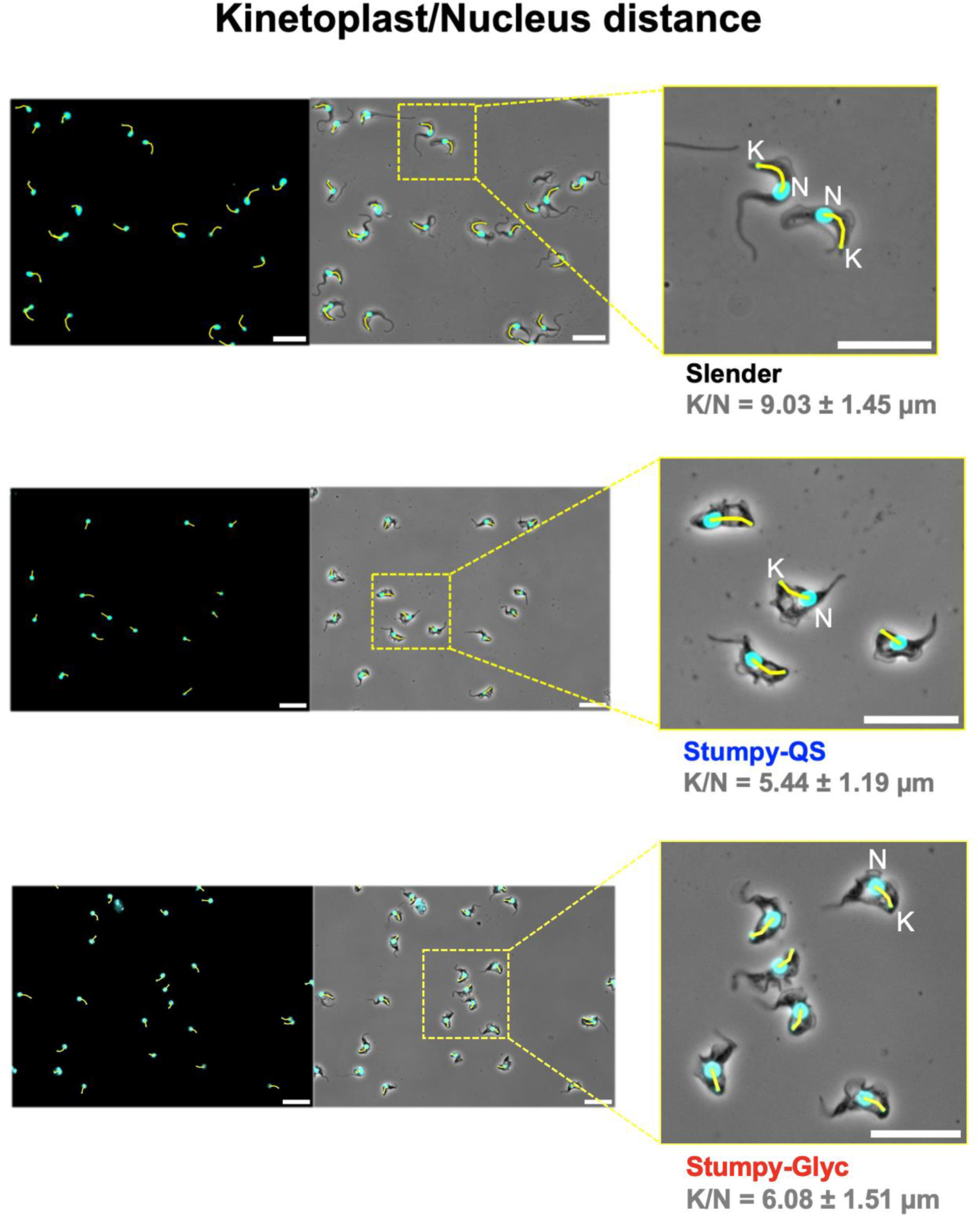
Stumpy-Glyc cells present a stumpy morphology. Example PC Phase contrast images of slender, stumpy-QS and stumpy-Glyc stained with DAPI for nucleus (N) and kinetoplast (K) localization. Yellow lines indicate how the distance between the nucleus and kinetoplast was measured manually using ImageJ software. White scale bars = 20 µm.

**Figure S4.**
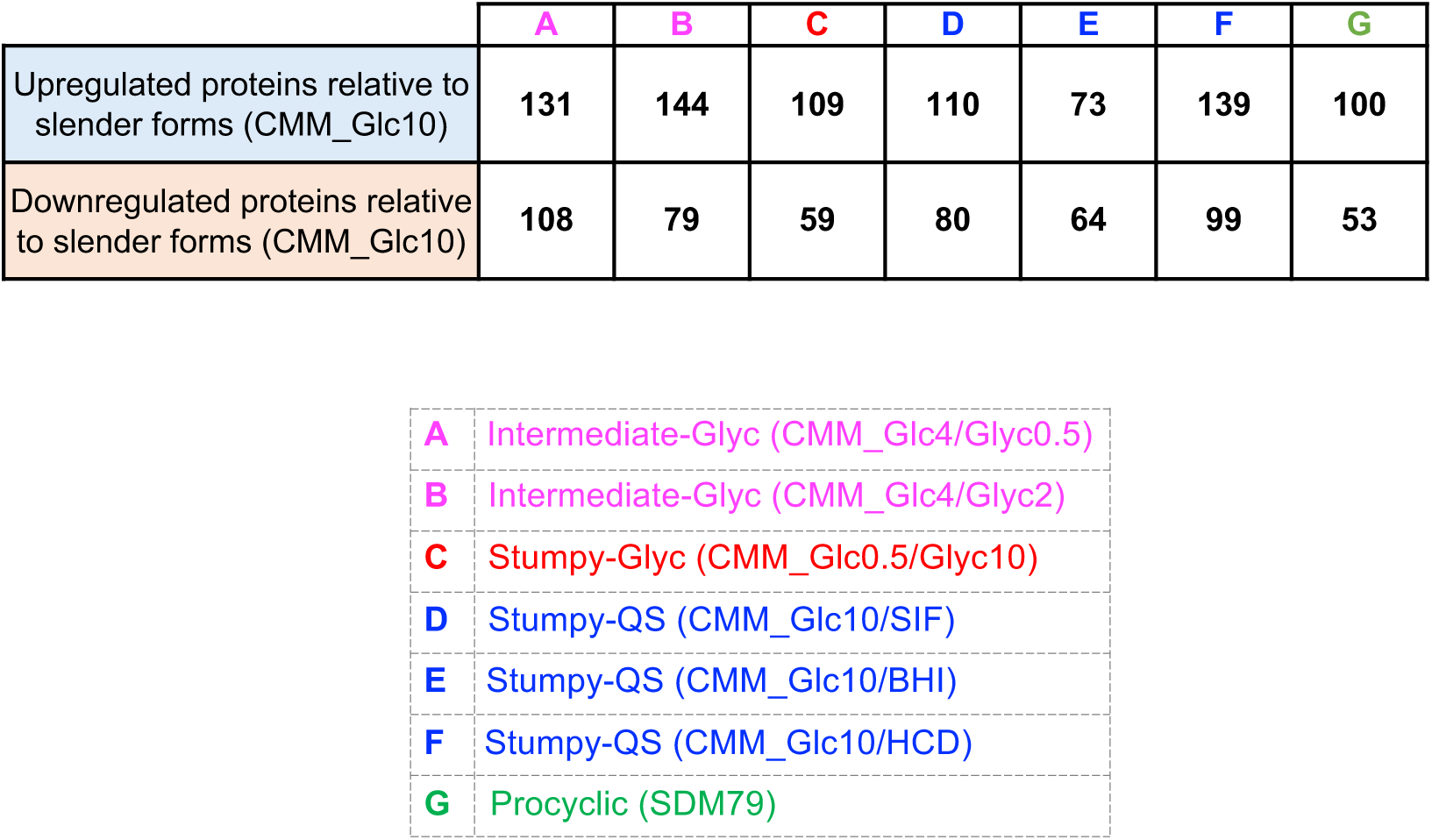
Number of differentially regulated proteins across *T. brucei* life cycle stages relative to slender forms (CMM_Glc10) This table displays the numbers of differentially upregulated and downregulated proteins— across various *T. brucei* life cycle stages—relative to slender forms (CMM_Glc10). The conditions include “Intermediate-Glyc” (CMM_Glc4/Glyc0.5 and CMM_Glc4/Glyc2), “Stumpy-Glyc” (CMM_Glc0.5/Glyc10), “Stumpy-QS” (CMM_Glc10/SIF, CMM_Glc10/BHI, CMM_Glc10/HCD), and “Procyclic” (SDM79). The names of the proteins regulated in each form, with their accession numbers, are provided in an Excel table.

**Figure S5.**
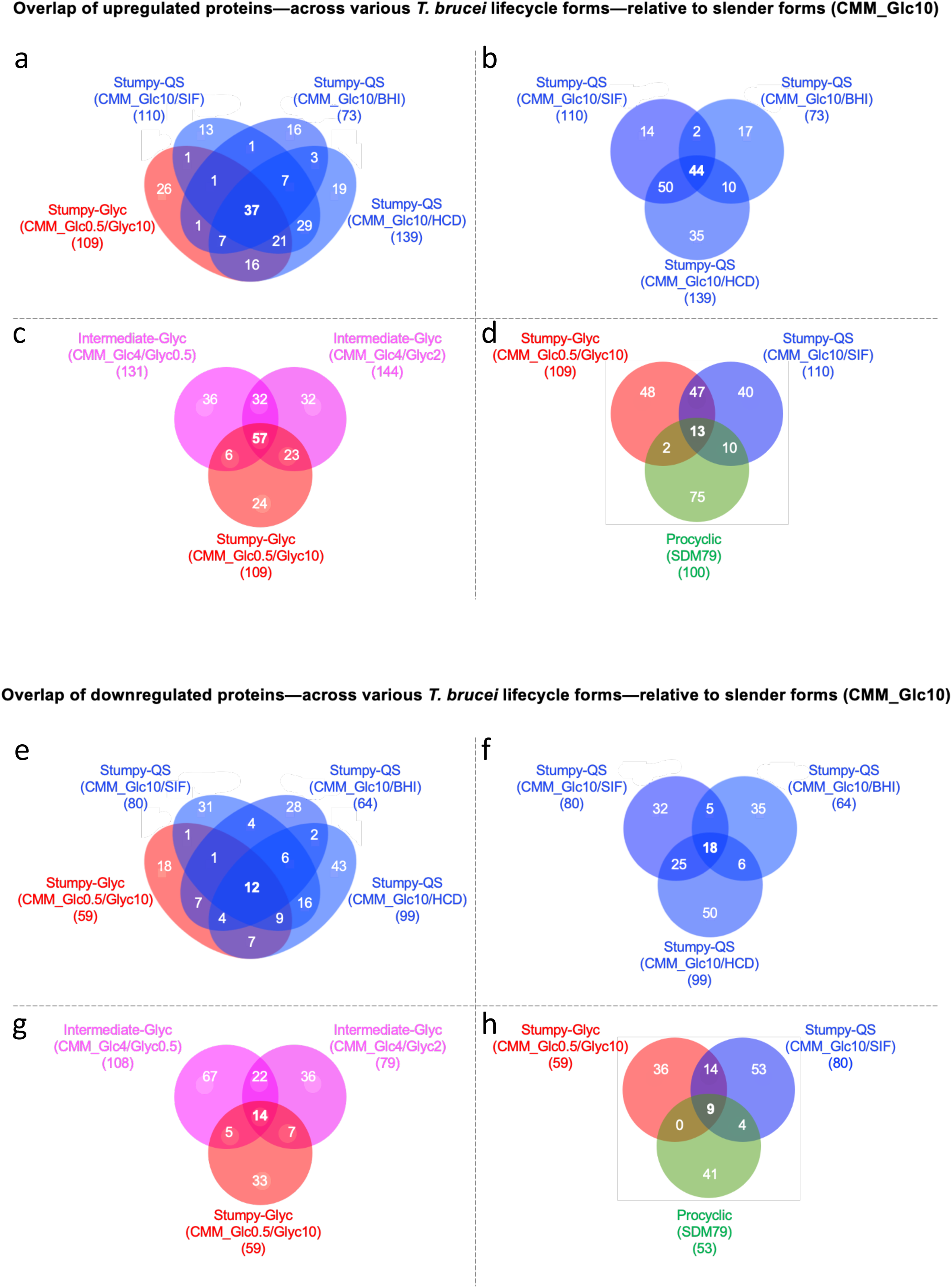
Stumpy-Glyc protein expression profile is closer to that of Stumpy-QS than to the PCF one. This figure presents a series of Venn diagrams that illustrate the overlap of differentially regulated proteins, both upregulated (upper panel) and downregulated (lower panel)—across various *Trypanosoma brucei* life cycle stages—relative to the slender forms. For clarity, “regulated proteins” refers to proteins that are differentially regulated relative to the slender form (CMM_Glc10). The number of regulated proteins relative to slender forms is represented for each condition under the name in parentheses. **a,** compares these upregulated proteins in the “Stumpy-Glyc” condition (CMM_Glc0.5/Glyc10) (red) with those in three “Stumpy-QS” conditions (blue): CMM_Glc10/SIF, CMM_Glc10/BHI, and CMM_Glc10/HCD. In this comparison, 37 proteins are consistently upregulated across all Stumpy-QS and Stumpy-Glyc conditions. **b,** shows the overlap of upregulated proteins among Stumpy-QS conditions alone, identifying 44 proteins commonly upregulated relative to slender forms. **c,** illustrates the overlap in upregulated proteins between the “Stumpy-Glyc” condition and two “Intermediate-Glyc” conditions (pink), revealing 57 proteins upregulated across all these conditions. **d,** highlights the overlap of upregulated proteins across “Stumpy-Glyc” (red), SIF-induced “Stumpy-QS” (blue), and “Procyclic” (SDM79) (green) conditions, with only 13 proteins upregulated in all three. **e-h**, mirrors the upper panel but displays downregulated proteins instead of upregulated, relative to the slender forms.

**Figure S6.**
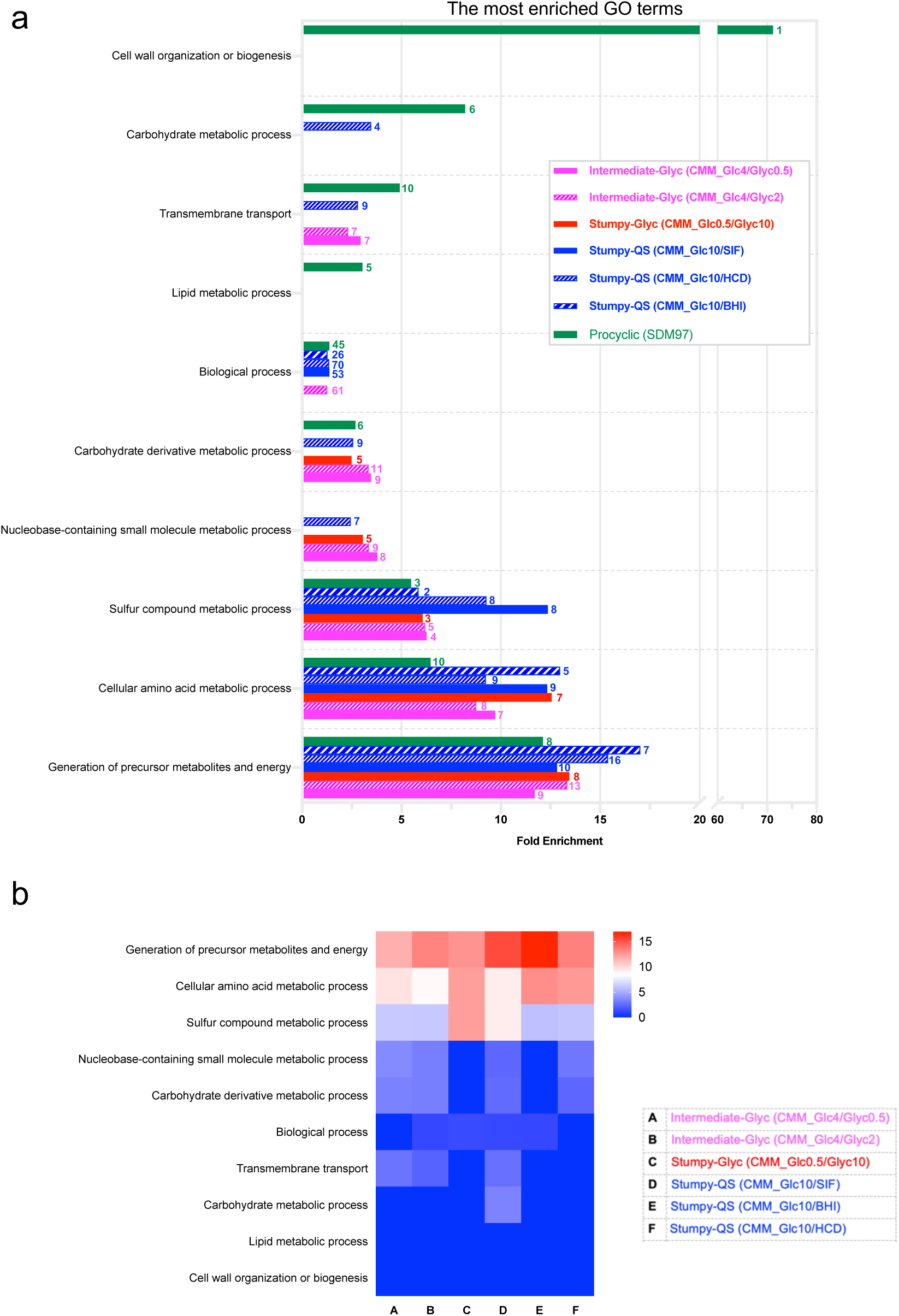
GO term enrichment analysis for biological processes of genes upregulated by >2 folds in different conditions as compared to slender forms (CMM_Glc10). **a**, shows terms on the y-axis that are significantly enriched (p<0.05) by more than two folds, comparing procyclic, stumpy-Glyc, stumpy-QS, and intermediate-Glyc forms. The numbers at the right end of the bars indicate the number of genes specific to each GO term. **b**, shows a heat map showing the enriched GO terms in different conditions compared to slender forms.

**Figure S7.**
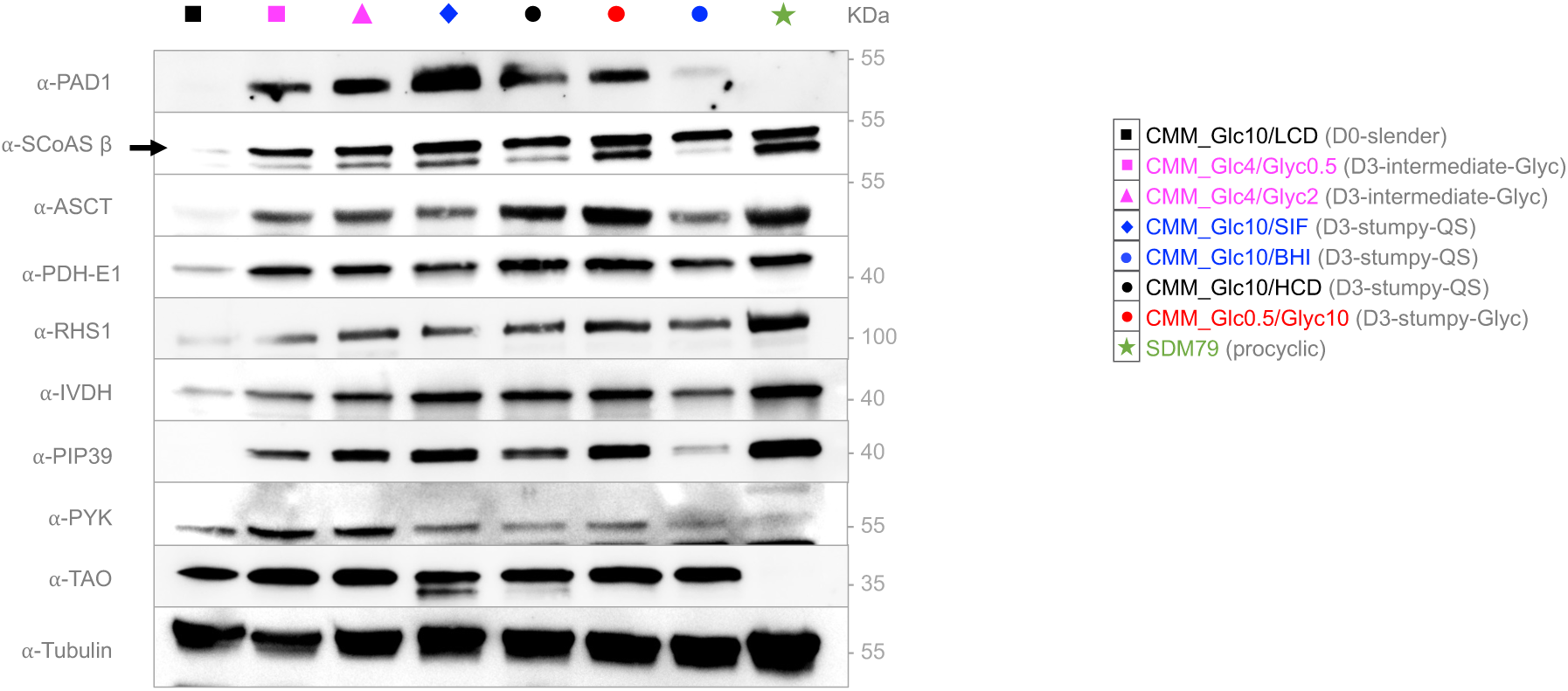
Stumpy-Glyc cells have a similar protein expression pattern as stumpy-QS. Western blot analysis confirming the proteomics data using the different forms analyzed in the proteomics study, with a range of available antibodies.

**Figure S8.**
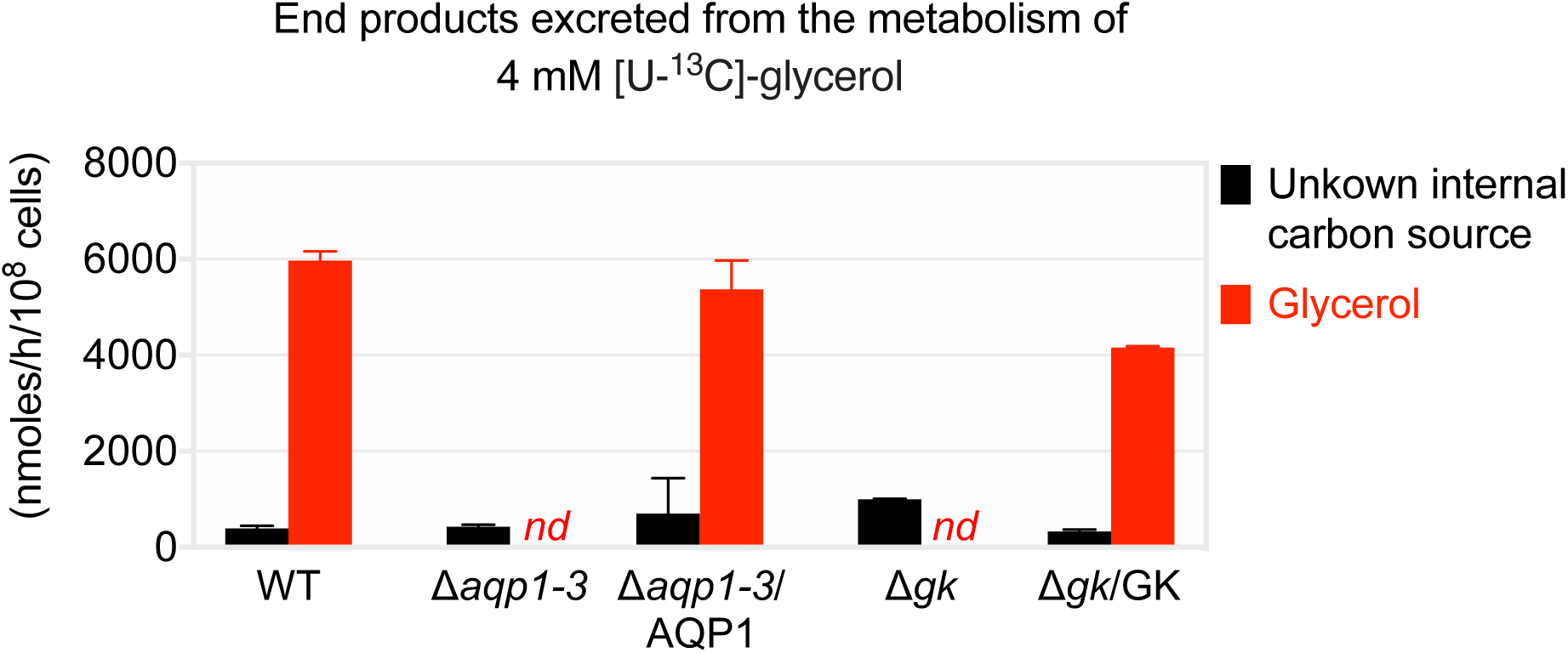
Validation of glycerol kinase and aquaglyceroporin null mutants and the corresponding rescue cell lines by ^1^H-NMR analysis. The parental (WT), knockout (Δ*aqp1-3* and Δ*gk*) and rescue (Δ*aqp1-3*/AQP1 and Δ*gk*/GK) cell lines were incubated for 1 h in PBS/NaHCO_3_ buffer containing 4 mM of uniformly ^13^C-enriched glycerol ([U-^13^C]-glycerol). The amounts of non-enriched and ^13^C-enriched metabolic end products excreted in the spent medium from the metabolism of an unknown internal carbon source (Bringaud et al., 2015) and [U-^13^C]-glycerol, respectively, were determined by ^1^H-NMR spectroscopy. *nd*, not detectable.

**Figure S9.**
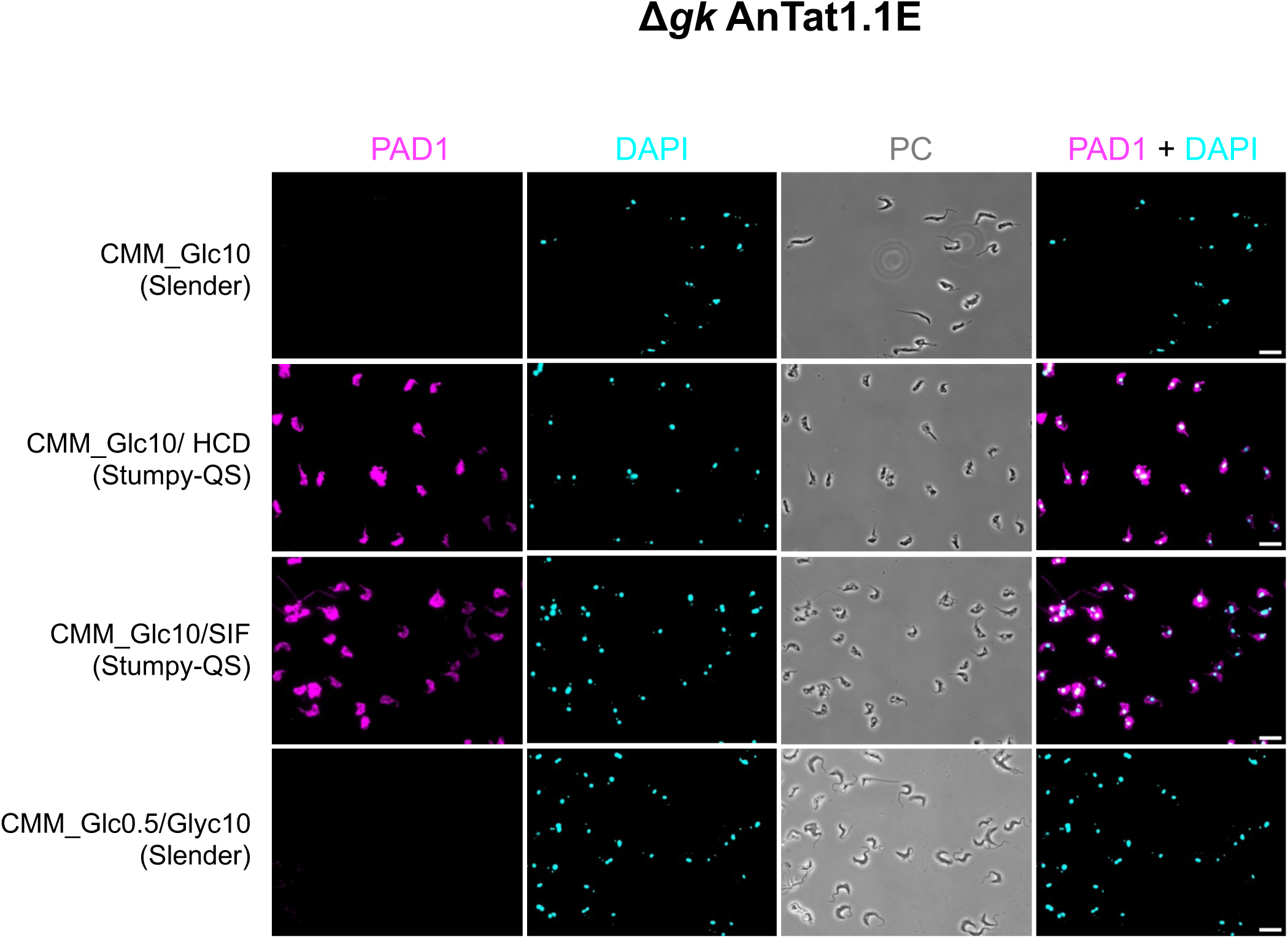
Glycerol kinase null mutant fails to differentiate to stumpy-Glyc when incubated with glycerol. Immunofluorescence analysis of the glycerol kinase null mutant (△*gk* AnTat1.1E) incubated in (*i*) CMM_Glc10 and maintained as slender forms in CMM_Glc10 by daily dilution to prevent at high cell density, (*ii*) in CMM_Glc10 at high cell density without daily dilution (CMM_Glc10/HCD), (*iii*) in SIF-rich CMM_Glc10 spent medium derived from a stationary phase culture of AnTat1.1E-GFP::PAD1_UTR_ (CMM_Glc10/SIF), and (*iv*) in low glucose (0.5 mM) CMM containing 10 mM glycerol (CMM_Glc0.5/Glyc10). PAD1 expression is revealed using anti-PAD1 (false colored magenta), nucleus and kinetoplast using DAPI (false colored cyan). PC Phase contrast. (n = 3 independent experiments). Scale bar = 20 µm.

**Figure S10.**
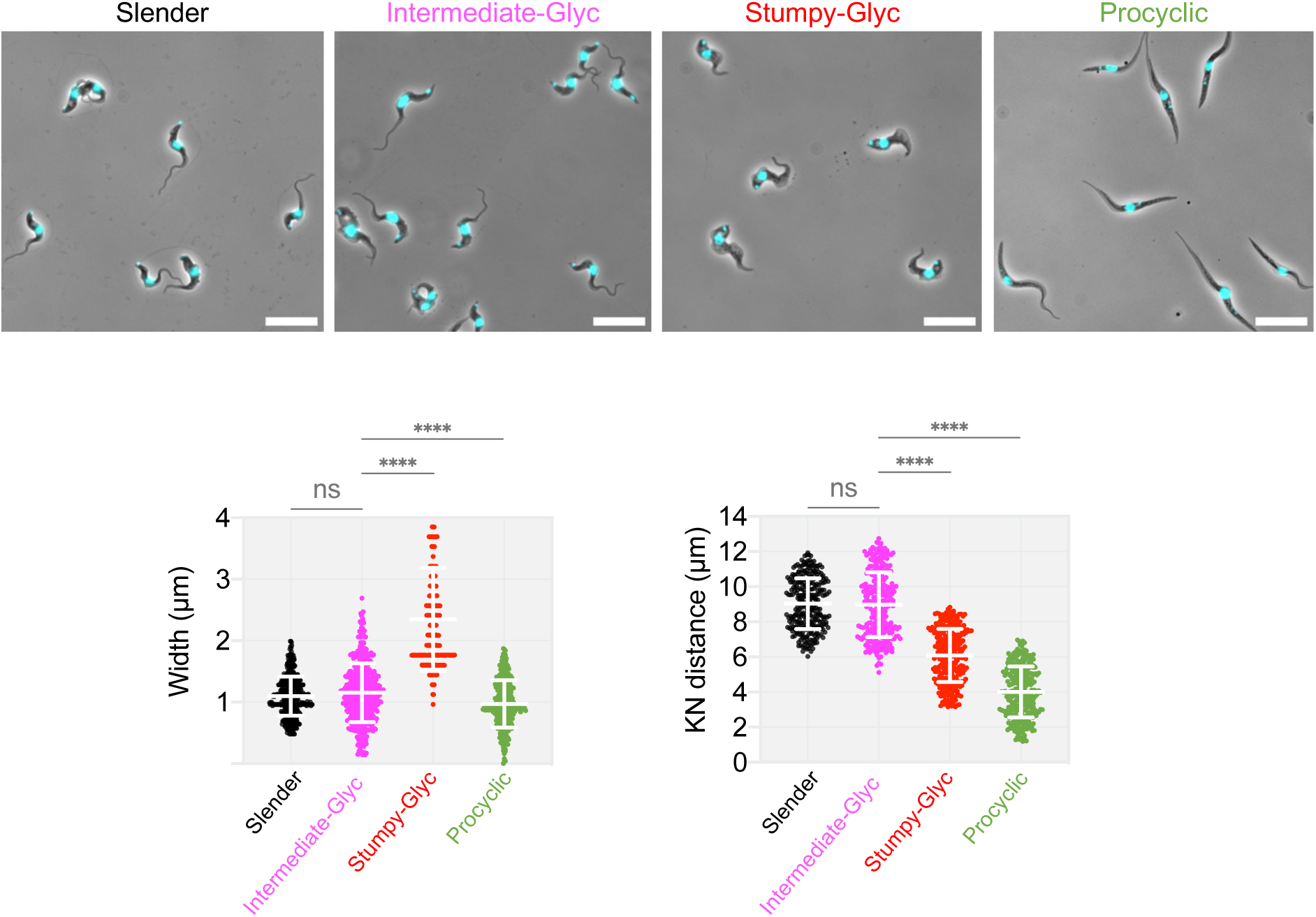
Intermediate-Glyc forms are slender-like forms. Phase contrast microscopy images overlaid with DAPI fluorescence, highlighting the nucleus and kinetoplast (false-colored cyan) of distinct morphological forms: Slender, Intermediate-Glyc, Stumpy-Glyc, and Procyclic forms. Scale bar = 20 µm. Intermediate-Glyc morphology was confirmed by measuring the cell width and the distance between the kinetoplast and the nucleus (K-N) of 250-300 cells per group, using a previously described ImageJ macro (****, p < 0.0001; ; ns, not significant).

